# Short steps can take you far: Phylogenetic analysis of Australasian *Cheilanthes distans* reveals frequent shorter-range dispersal

**DOI:** 10.1101/2024.06.22.600207

**Authors:** Karla Sosa

## Abstract

Biologists have long pondered species’ geographical distributions and sought to understand what factors drive dispersal and determine species ranges. In considering plant species with large ranges, a question that has remained underexplored is whether large ranges are attained primarily through many instances of short scale dispersal or whether instead widespread ranges are attained by propagules with increased dispersal distances. Ferns provide an ideal system to explore this question as their propagules are very small spores, which have been theorised can be carried by wind to essentially anywhere on the planet. Unfortunately, population-level genetic data in ferns is relatively uncommon, limiting our ability to answer this and related questions. For this work, I focus on *Cheilanthes distans* (Pteridaceae) as a study system, a widespread fern with extensive spore variation that occurs over Australia and into New Zealand/Aotearoa, New Caledonia, and other Pacific islands. I sampled widely across the species’ range, in addition to across Australasian *Cheilanthes* (as a robust tree for the genus does not exist), ultimately building a phylogeny based on the GoFlag 451 bait set. With these data, we can investigate additional questions, including whether reproductive mode, polyploidy, or lineage influence dispersal, as well as whether movement is occurring randomly or is instead asymmetrical. I explored the relationships between sexual and apomictic specimens to understand whether the former are the parental lineages to apomictic plants and whether we find evidence for apomictic plants dispersing out of a small parental range. I investigated how many times polyploid lineages have arisen in *C. distans* and whether they are each limited geographically, perhaps forming isolated ranges that collectively result in *C. distans’* larger range. Additionally, I generated estimates for ancestral ranges and dispersal between populations to understand whether certain lineages are limited to particular geographic regions, to explore the directionality of dispersal, and to assess whether most movement is happening over short or long distances. Particularly interestingly, I find that most dispersal in this species appears to occur over smaller steps rather than longer jumps, underscoring how short movement can nevertheless allow for establishment of large ranges; this dispersal is not limited phylogenetically and seems to occur equally for all lineages. What is more, I find evidence for asymmetrical dispersal directionality, apparently most frequently tracking trade winds. These findings demonstrate the importance of population-level data, and provide concrete results that add nuance to long-standing dispersibility hypotheses in the fern community that have, until now, lacked empirical data.

## 1. Introduction

Biologists have long pondered species’ geographical distributions and sought to understand what factors drive dispersal and determine species’ ranges (Linnaeus 1781; Von Humboldt and Bonpland 1805; Darwin 1859; Wallace 1876; Wilson and MacArthur 1967). For organisms that are mostly immobile—such as plants—species require propagules that will disperse and establish in new territory in order to expand their range. Several studies have shown that most plant propagules tend to land close to their parent plant, with only a few dispersing larger distances (Norros et al. 2014; Muller-Landau et al. 2008; Wilkinson et al. 2012 and citations therein). Although some of the species studied are widely distributed, these studies have looked at propagule dispersal only over small scales (up to 15km). To my knowledge, however, there have been no studies that look at the prevalence of longer-distance dispersal in more widespread species, nor any that assess its potential role in species attaining larger ranges. A question, then, remains to be explored: Are large ranges attained primarily through short scale dispersal, or could their widespread ranges be aided by propagules with increased dispersal distances?

Ferns provide an ideal system to study this question. The propagules of ferns are very small spores (∼40–80µm) produced in vast quantities (Moran 2004). Because of this, it has been theorised that fern spores can be carried by wind to essentially anywhere on the planet (Tryon 1970; Wolf et al. 2001). In fact, the elevated proportion of ferns on oceanic islands has been used as evidence of their dispersibility (Tryon 1970; Smith 1972; Peck et al. 1990). What is more, many fern species are in fact widely distributed (Smith 1972), meaning dispersal must be frequent in this group. Several little-explored questions follow: Is most of the dispersal in ferns in fact happening over long ranges, or is most of it occurring over short hops that ultimately lead to wide distributions? Is dispersal influenced by reproductive mode? Are certain lineages better dispersers than others? And do ferns disperse widely and randomly, or is movement asymmetrical? For example, a handful of studies available hint at dispersal in plant lineages that coincides with asymmetrical dispersal driven by trade wind direction (Muñoz et al. 2004; Sanmartín et al. 2007). However, there has been surprisingly little population-level data collected in ferns (Pelosi and Sessa 2021), which is necessary to understanding dispersal dynamics within a species.

To explore these questions, I focus on *Cheilanthes distans* (Pteridaceae) as a study system, a widespread fern that occurs over Australia and into New Zealand/Aotearoa, New Caledonia, and other Pacific islands (Tindale and Roy 2002). This species is primarily composed of apomictic polyploids, but a few sexual specimens are known that inhabit a narrow range in northeastern Australia. I have also observed extensive variation in spore size in this species, a trait which seems to influence dispersal (Sosa 2024). I thus wished to gain an understanding of how reproductive mode, spore size, and phylogeny are influencing dispersal and biogeography of *C. distans*. Because no phylogenetic tree exists for Australasian *Cheilanthes* as a whole, it was necessary to first construct a robust tree that could be used for further analyses. Given known polyploidy in the genus (Tindale and Roy 2002), as well as frequent issues of discordance in related groups (Dyer et al. 2012; T-T. Kao, pers. comm.), I took advantage of recent advances in high-throughput sequencing and utilised sequence capture techniques (Weitemier et al. 2014) and the flagellate plant GoFlag 451 bait set (Breinholt et al. 2021) to collect sequence data over hundreds of single-copy nuclear loci. As my focal interest is on *C. distans*, I included extensive sampling for this species in particular in order to build a population-level phylogeny.

With a phylogeny, I then explore the relationships between sexual and apomictic specimens to understand whether the former are the parental lineages to apomictic plants and whether we find evidence for apomictic plants dispersing out of a small parental range. I also investigate how many times polyploid lineages have arisen in *C. distans*, whether they are each limited geographically (perhaps forming isolated ranges that collectively result in *C. distans’* larger range), and what the relationship of polyploidy is to reproductive mode. Considering findings that plants with medium-sized spores have the largest ranges (Sosa 2024), with the phylogeny I also ask whether spore size shows phylogenetic signal within *C. distans*, and use the sequence data to investigate genetic diversity among spore size groups to search for additional evidence of wide dispersal. Finally, I generate estimates for ancestral ranges and dispersal between populations to understand whether certain lineages are limited to particular geographic regions, the directionality of dispersal, and whether most movement is happening over short or long distances.

I was able to build a robust phylogeny for nearly all Australasian *Cheilanthes*, with high recovery across most loci (including non-targeted chloroplast loci), and found that taxonomic breadth is more important than sequence completeness in generating a robust tree. My analyses then revealed intriguing patterns for the relationship between presumed sexual specimens of *C. distans* and their apomictic relatives: while it is possible for the sexual specimens to be parental, the phylogeny does not immediately lead to this conclusion. Particularly interesting as well, I found that most dispersal in this species occurs over smaller steps rather than longer jumps; this dispersal is not limited phylogenetically and seems to occur equally for all lineages. What is more, I find evidence that dispersal direction appears asymmetrical and may most frequently be tracking trade winds. These findings provide concrete results that add nuance to long-standing dispersibility hypotheses in the fern community that have, until now, lacked empirical data.

## 2. Materials and Methods

### 2.1 Taxon sampling

In order to build a phylogeny that would place the *Cheilanthes distans* specimens into phylogenetic context, specimen selection aimed to sample all accepted *Cheilanthes* species found in Australia as per Chambers and Farrant (1998), which is the most recent review of this genus in the region. Accepted species—besides *Cheilanthes distans*—include: *C. adiantoides, C. austrotenuifolia, C. brownii, C. caudata, C. contigua, C. fragillima, C. lasiophylla, C. nitida, C. nudiuscula, C. praetermissa, C. prenticei, C. pumilio, C. sieberi,* and *C. tenuifolia*. Of these, I was able to obtain material for all but *C. austrotenuifolia*. Additional specimens that presented intermediate or distinct morphologies were also sampled. Among these, I included a couple indeterminate species found during my fieldwork. I also included a couple of ‘contentious’ specimens, in particular a specimen noted as *C. cavernicola* by Jones (1988), which is stated by him to be related to *C. tenuissima* (a species described by Quirk et al. (1983)); however, *Cheilanthes cavernicola* is subsumed by Chambers and Farrant (1998) into *C. pumilio*, while *C. tenuissima* is subsumed into *C. caudata*. I also included a specimen noted as *C. shirleyana* by Quirk et al. (1983), a species which is subsumed by Chambers and Farrant (1998) into *C. tenuifolia*. I endeavoured to include two samples per accepted species when possible, the sole exception being *C. tenuifolia* for which I included wider sampling that will aid in studying spore variation in this species at a later date (Sosa, unpublished).

For the sampling of apomictic *Cheilanthes distans*, I aimed to include as wide a geographic range as possible from those samples that had been studied in Sosa (2024), in addition to sampling as widely as possible among spore sizes. I also included all specimens identified as presumed sexual plants. To create an outgroup, I included specimens of *Cheilanthes ecuadorensis, C. incarum, C. micropteris, C. obducta, C. pilosa,* and *C. rufopunctata*; these species are all South American and were found to be earlier diverging and sister to all Australasian *Cheilanthes* (Sosa et al. 2021). For all samples, I preferentially sampled silica dried material when available, and otherwise sampled herbarium material. All samples included are listed in SI1. Amends and apologies are owed to the Traditional Custodians and Indigenous Nations from whose lands these samples were taken without consent nor compensation.

### 2.2 DNA EXTRACTION AND SEQUENCING

For DNA extractions, approximately 1cm^2^ of leaf material was sampled from each specimen, either from herbarium sheets or silica-dried leaf material. Isolations were carried out using Omega EZNA SP Plant DNA Kits with standard protocols, as this has shown good results in other *Cheilanthes* species (L. Huiet, pers. comm.). Because many of the samples were derived from herbarium specimens, first elutions were used for sequencing since these tend to have higher concentrations of genetic material. Extractions were packaged in dry ice and shipped overnight from Durham, NC to collaborators in Gainesville, FL, and kept frozen until delivered to RAPiD Genomics (Gainesville, FL). Extractions were screened for DNA concentration by RAPiD Genomics; any sample found to have 0ng of DNA was eliminated from the sequencing pool. The remaining samples were then sequenced using the GoFlag 451 target enrichment probes designed by Breinholt et al. (2021) to capture exons in 409 single- and low-copy nuclear genes in flagellate plants (a key aspect considering frequent polyploidy amongst Australasian *Cheilanthes*), and following the methods described therein. Raw sequence data used for this project can be accessed at the following repository: https://doi.org/10.7910/DVN/BL7FMR. Additional sequence data that I was unable to include in the analyses below due to delays in sequencing can be found at the following repository: https://doi.org/10.7910/DVN/XJW1XE – please use it!

### 2.3 Sequence filtering and data sets

Raw sequence reads were cleaned for very short fragments (< 35bp) and the ends trimmed (3bp on either end) using Trimmomatic (Bolger et al. 2014); adapters and barcodes had already been removed by RAPiD. Resulting sequences were then filtered using HybPiper (Johnson et al. 2016). HybPiper requires target sequences to which to align raw reads. Target sequences were assembled by concatenating all the reference genes used for bait design by Breinholt et al. (2021). Although chloroplast sequencing is not targeted by the baits, the large numbers of chloroplasts present in a cell mean that there is often ‘spill-over’ sequencing of plastomes (e.g. Schneider et al. 2021). For this reason, I also included the chloroplast sequences of *Cheilanthes micropteris* generated by Robison et al. (2018) in the reference file. All target sequences used can be found in the following repository: https://doi.org/10.7910/DVN/1TGMRV. In order to visualise sequence recovery over all loci (including chloroplast regions), a heatmap was generated using the script provided within HybPiper and executed in R version 3.6.0 (this version of R also used for all subsequent analyses; R Core Team 2017).

As mentioned above, polyploidy is frequent in this group, and homologous loci are necessary to ensure accurate phylogenies. In order to resolve potential paralogy I used paralog_investigator within HybPiper to first identify the number of paralogs for each locus. I removed any loci with more than 10 paralogs from the study; while these are not necessarily sequencing errors and may instead be due to the limits of assembling 100bp fragments, it becomes very difficult if not impossible to disentangle homology at such high numbers. I then built paralog trees for the remaining loci using the automatic alignment option in mafft (Katoh and Standley 2013), and the GTR model to generate approximate-maximum-likelihood trees in FastTree (Price et al. 2010). I reviewed all trees for structure among paralogs. Among the trees, I observed three patterns (for examples of these see SI2 – Fig. 1): (1) trees that showed clear structure signalling locus duplication, (2) trees where all paralogs clustered into a monophyletic group, and (3) trees where paralogs showed no clear structure. For trees of type (1), I ensured that the sequence copy retained for downstream analyses was that which clustered with the majority of the samples (since some of these are diploid and thus only appear once). For trees of type (2), I made no changes, leaving the sequence copy that had been selected by HybPiper (based on higher coverage and similarity to target loci) for downstream analyses. I eliminated any loci that formed trees of type (3), since it is not possible to determine homology under these circumstances.

**Figure 1:**
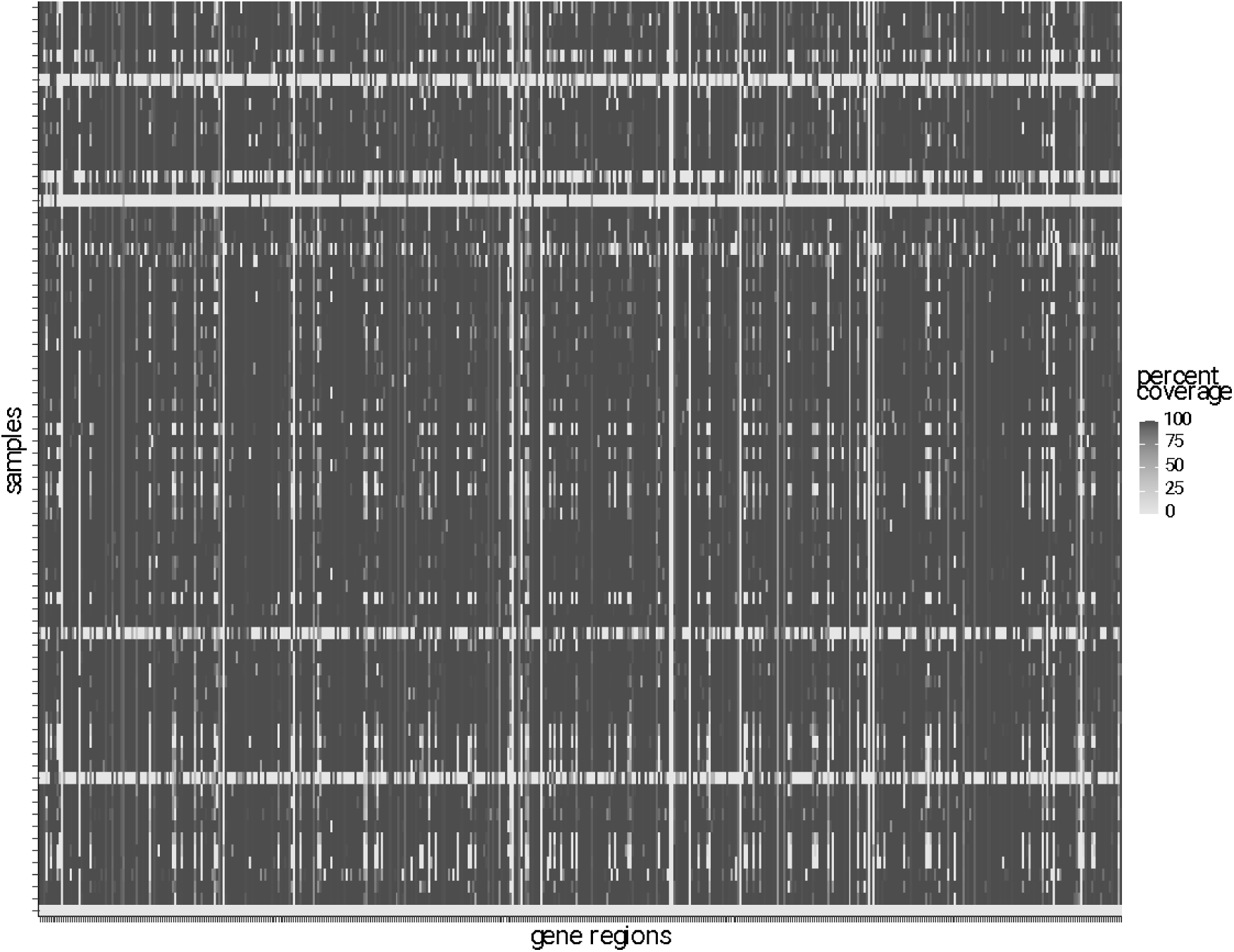
Heatmap showing the percent coverage of recovered sequences for each sample (row) and each locus (column), these loci including chloroplast regions.

Workers over the years have explored whether phylogenies are calculated most accurately with fewer, higher coverage sequences vs. more, lower coverage sequences (Heath et al. 2008; Poe and Swofford 1999; Hillis et al. 2003). While most of our recent data suggest phylogeny inference is best served by having as much data as possible (Zwickl and Hillis 2002; e.g. Pollock et al. 2002), because these studies were done on data that are considerably different from my own, I decided to assemble three data sets with different levels of stringency. The least stringent data set included loci with sequences at least 10% of the length of the original locus, samples with at least 10% of loci present and loci with at least 10% sample coverage (these last two calculated after the 10% minimum length filtering had been applied). The most stringent data set filtered minimum sequence lengths at 90% coverage, samples at 90% of loci present, and loci at 90% of samples present. The intermediate data set used a cut-off of 50% minimum sequence length, 50% of loci present per sample, and 50% of samples per locus. To assemble these data sets, I calculated sequence lengths (get_seq_lengths in HybPiper) for all loci that remained after paralog filtering. I then extracted sequence fasta files that included introns beyond the exon-only baits (intronerate in HybPiper), and created three separate folders that contained loci for each of the 10%, 50%, and 90% stringency levels.

### 2.4 Phylogenetic analyses

Alignments were done separately for each locus using MUSCLE (Edgar 2004). The relevant alignments for each analysis were concatenated into super matrices using SuperMatrix in evobiR (Blackmon et al. 2013) in R. All final alignments for each of the stringency levels described above can be found at the following repository: https://doi.org/10.7910/DVN/AUHWGP; the code used for Methods Sections 2.3–5 can also be found within this repository, as can the data used for Section 2.5 below.

I first estimated phylogenies for each of the stringency levels for chloroplast and nuclear sequences separately, using both maximum likelihood and Bayesian inference methods. I estimated maximum likelihood trees with RAxML (Stamatakis 2006) using the GTR+gamma model on concatenated sequences with a partition per locus, and bootstrap number set to autoMRE (i.e. the extended majority-rule consensus tree criterion). I estimated Bayesian trees using MrBayes (Ronquist and Huelsenbeck 2003) with a single partition given the complexity of the data, using a beta transition/transversion rate ratio and a Dirichlet substitution rate for the GTR model, with uniform priors for gamma and proportion of invariant sites. I ran Bayesian analyses for 50 million generations, with two runs and 8 chains and a 0.25 burn-in. Once I corroborated there was no significant discordance between nuclear and chloroplast trees within stringency data sets within methods, I re-ran analyses with combined nuclear and chloroplast data (unpartitioned for MrBayes and partitioned for RAxML). Additionally, I used coalescent methodology to generate a species tree estimate. I first estimated individual trees for all nuclear loci using RAxML with a GTR+gamma model, and used these gene trees to estimate a species tree with ASTRAL (Mirarab and Warnow 2015). All RAxML and ASTRAL analyses were carried out on the Duke University Compute Cluster, while MrBayes analyses were run on the CIPRES Science Gateway (Miller et al. 2011).

I compared the final trees for each of the analyses (RAxML, MrBayes, ASTRAL) across each of the stringency levels (10%, 50%, 90%), as well as comparing the three stringency levels within a single method. I assessed whether there were significant areas of discordance amongst trees, whether any taxa or groups seemed to ‘jump around’ the tree, and how the support values varied from tree to tree, ultimately selecting a single ‘best tree’ from among these nine options (MrBayes at 10% stringency).

### 2.5 Ploidy estimation

I estimated ploidy for all samples using nQuire (Weiß et al. 2018), executed in an Amazon LightSail Linux virtual machine. This method uses alignments to make SNP calls for each specimen, from which distributions of SNP proportions are then calculated. In other words, SNP calls are plotted on a histogram to then determine ploidy: diploids should show a normal curve with a peak centred at a 1:1 ratio of SNPs (50%); triploids should show two normal curves with peaks at 33% (1:2) and 66% (2:1); and tetraploids should show all of the above, with peaks at 33%, 50%, and 66%. This programme has been shown to be accurate on herbarium specimens and with sequence capture data (Viruel et al. 2019). To carry out my analyses, I determined loci for which HybPiper had detected at most four paralogs (see 2.3 for details on paralog detection); while samples with higher ploidies might be possible, nQuire is only able to accurately detect ploidies up to 4x. If a specimen with higher ploidy existed it should still be detectable and would result in an inconclusive estimate, thus ultimately being excluded from the study; ultimately this cut-off results in a conservative dataset. I created a reference ‘genome’ made of the above selected loci, concatenating the relevant reference genes used in bait design for the most closely related species to *Cheilanthes*, giving preference to *Gaga arizonica* sequences if available and otherwise using *Notholaena montieliae*; this reference ‘genome’ is the same as the one used in the methods of Sosa (2024). Raw sequence reads for all samples were then aligned to the reference ‘genome’ using bwa (Li 2013), and sorted using SAMtools (Li et al. 2009). These BAM were the files then uploaded into nQuire, denoised, and ploidy estimations calculated.

### 2.6 Exploring reproductive mode and ploidy

In ferns, sexual diploids are widely considered to be the progenitors to apomictic polyploid lineages (Grusz et al. 2021 and references therein). One of the aims of this study was to test whether the presumed sexual specimens identified in Sosa (2024) were in fact parental lineages to apomictic *C. distans*. A related aim was to generate an initial estimate for how many times polyploidy has arisen in this species. Especially given that I found both triploid and tetraploid lineages, their phylogenetic relationships to each other is also of interest. To visualise patterns in reproductive mode, I plotted states onto the tips of the tree, then reconstructed ancestral states for reproductive mode using a maximum likelihood framework and testing models with transitions having either ‘equal rates’ or ‘all rates different’. I plotted the best fitting model onto the nodes of the tree in order to trace state changes across the phylogeny. To further understand where changes in state are happening along the tree, I also generated stochastic character maps—which sample character histories from their posterior probability distribution using an MCMC approach—sampling 100 trees using the best fitting model, and plotted the character changes estimated across all 100 trees onto a single phylogeny. All analyses were carried out using phytools (Revell 2012) and all plotting was done with ape (Paradis et al. 2004), both executed in R. To study patterns in ploidy, I repeated the above analyses, replacing the reproductive mode character data with the ploidy data.

To further understand how these traits may be changing over the tree, and because certain models of evolution can’t be implemented during ancestral state reconstruction, I built models for character evolution for both reproductive mode and ploidy using geiger (Pennell et al. 2014) in R. For reproductive mode I tested the following transition rate models: ‘equal rates’, ‘all rates different’, and irreversible models from sexual to apomictic, and from apomictic to sexual. For ploidy I tested the following transition rate models: ‘equal rates’, ‘all rates different’, meristic (meaning ploidy changes occur from 2 to 3 to 4 to 3, in other words allowing for ordered reversals), and ordered (meaning ploidy changes can only occur from 2 to 3 to 4 with no reversals). I calculated the weighted AIC scores for all models (which standardise AIC scores of alternative models in order to measure the relative weight of evidence for each model) to determine the best fitting model.

To test for phylogenetic signal of reproductive mode, I calculated the Fritz and Purvis’s D statistic (Fritz and Purvis 2010)—which measures the number of sister-clade differences—using caper (Orme et al. 2013) with 10,000 iterations in R. A value of 0 means the trait is evolving under Brownian motion, while a value of 1 means the trait is evolving randomly (i.e. there is no phylogenetic signal); a value less than 0 would signal a highly conserved trait, and a value greater than 1 signals an over-dispersed trait. Because this test can only be done on binary data, I did not test for phylogenetic signal of ploidy.

It is frequently observed that shifts to apomixis are specifically linked with polyploidisation (Hörandl 2006). To test this hypothesis in my system, I ran a phylogenetic generalised least squares (PGLS) test between ploidy and reproductive mode (model = reproductive mode ∼ ploidy) using nlme (Pinheiro et al. 2007) in R; this method takes the phylogeny into account when estimating correlations between traits. I visualised the distribution of ploidy to reproductive mode using a mosaic plot.

### 2.7 Geographic patterns across the phylogeny

Given the finding that medium-sized spores appear to attain the largest ranges (explored in Sosa 2024), it is of interest as to how these dynamics translate to the phylogenetic level: do we see that medium-sized spores are over dispersed across the phylogeny, meaning they encompass greater genetic diversity? Or are certain lineages tending towards producing spores of certain sizes? I tested for phylogenetic signal of spore size in *C. distans* using two metrics: Blomberg’s K (Blomberg et al. 2003)—which compares the variance of phylogenetically independent contrasts to their expectation under Brownian motion—and Pagel’s lambda (Pagel 1999)—which stretches tip branches relative to internal branches depending on trait distribution; both of these were implemented in phytools (Revell 2012). A K value of 1 means relatives resemble one another as much as we expect under Brownian motion, while values less than one mean there is less phylogenetic signal than expected, and the inverse for values greater than one. Meanwhile, a lambda value of 0 means the phylogenetic signal observed is that expected from a star phylogeny (in other words, there is no phylogenetic signal), while a lambda of 1 means the signal matches the original tree. These metrics should inform us as to whether spore size is distributed randomly across the *C. distans* tree, whether it is highly conserved within lineages, or whether there is no signal beyond that expected as a result of phylogenetic relatedness.

If spores of certain sizes are in fact dispersing more readily, we might possibly expect their genetic diversity to be greater. This could be the case if some amount of sexual reproduction is occurring, even if very small (since we know that apomictic ferns still produce functional sperm (Walker 1962, 1985)), or alternatively if recurrent formations of apomictic lineages are transferring greater parental genetic variation with some also being more successful at dispersal if they have certain spore characteristics (i.e. medium size). In order to test whether certain spore size groups have higher genetic diversity, I estimated pi for all *C. distans* samples and tested for correlations between size and pi. For this, I first generated an invariant sites VCF based on the reduced dataset used for ploidy estimation (see 2.5) using mpileup in bcftools (Danecek et al. 2021). Samples were categorised into groups based on their mean spore sizes and using the same 20% splits (5 bins) used in Sosa (2024); given the small number of samples, to use any other smaller grouping would have resulted in singleton samples. I then used pixy (Korunes and Samuk 2021) to calculate pi for each of the groups, and finally generated a linear regression in R testing the effect of spore size groups, ploidy, and the interaction of spore size and ploidy on pi. Ploidy and the interacting effect were included as higher ploidies should on average exhibit higher genetic diversity (given more copies per gene), and as higher ploidy was found to be correlated with spore size (see results of Sosa 2024).

I also wished to explore whether lineages may be clustering geographically, as well as predicting the ancestral range for Australasian *Cheilanthes* and *C. distans* more specifically. Firstly, I categorised all samples into geographic regions, respecting areas of endemism in Australian flora (Crisp et al. 2001) and further defined by clusters observed on a map (SI2 – Fig. 5). However, the breaks between clusters varied in size and it is hard to say with certainty that a break is not due to a bias in sampling. For this reason I also categorised samples into a second set of larger geographic regions (Fig. 5), and carried out all analyses using both these groupings (i.e. smaller and larger regions). To understand dispersal between regions, I estimated models of evolution—using both ‘equal rates’ and ‘all rates different’ transitions between regions—for both large and small regions, and calculated weighted AIC scores to determine the best fitting model. In order to test for phylogenetic signal of geography (albeit roughly), and whether it was highly conserved (meaning that lineages are restricted geographically) or perhaps over dispersed (meaning that dispersal is more than we would expect by chance and therefore possibly adaptive), I also calculated both the K statistic and lambda using both latitude and longitude as proxies. To further elucidate geographic movement of species and lineages—if somewhat crudely—I followed the methods outlined in 2.6 and I plotted geographic character states onto tree tips, and then estimated ancestral states using both large and small geographic regions, testing both ‘equal rates’ and ‘all rates equal’ transitions between regions. I sampled 100 stochastic character maps—estimated using the best fitting model for both large and small regions—and mapped these changes onto the tree.

Because the topology of the tree will influence the final results of my analyses, and taking into account likely hybridization and/or incomplete lineage sorting observed in a subset of the tree different from my focal group (see 3.3), I repeated all analyses with a tree that comprised only *C. distans* samples. To further visualise the geographic distribution of *C. distans* lineages, I plotted its phylogeny directly onto a map using phylo.to.map from phytools.

## 3. Results

### 3.1 DNA EXTRACTION AND SEQUENCING

Of the 100 DNA extractions, 14 failed altogether (DNA concentrations = 0ng). Of the remaining 86 samples, eight of these failed sequencing. This resulted in a total of 78 sequenced specimens. I was able to obtain at least one sample per species with the exceptions of *C. praetermissa, C. prenticei,* and *C. shirleyana*. The failed samples were often those extracted from older herbarium material, although many herbarium samples were successful.

### 3.2 Sequence filtering and data sets

I used a total of 496 reference loci onto which to map sequencing reads; 87 accounted for chloroplast regions and 409 for the targeted nuclear loci. The sequence heatmap shows high recovery for most samples and across most gene regions (Fig. 1): the average number of loci recovered per sample was 454 (91.5%), with the number of loci with 75% sequence coverage averaging 423 (85.3%). Of the chloroplast loci—despite not being directly targeted—76 had gene recovery in over half the samples (87.4%), with two other loci only being recovered in 22 and 36 samples respectively.

I eliminated ten loci because their average paralog count was ten or higher. The remaining loci averaged 2.4 paralogs, with a maximum of seven paralogs and a minimum of zero. I visually inspected all trees that had paralogs above zero (a total of 382 trees). Of these, 11 required some swapping of the sequences that HybPiper had selected as the ‘main’ sequence for a different paralog because the ‘main’ sequence for these samples was not homologous to the ‘main’ sequence for most of the other samples (meaning the ‘main’ sequence selected was a paralog). Five of these trees required only one swap for a single sample, while the rest required more than one change.

The final filtered data sets comprised 74 taxa and 472 loci for 10% stringency, 72 taxa and 469 loci for 50% stringency, and 62 taxa and 417 loci for 90% stringency. The numbers of sequences, taxa, and loci removed at each level are displayed in Table 1.

**Table 1:** Total data removed by category for each of the minimum coverage stringency schemes.

| Minimum coverage | Sequences removed (due to insufficient length) | Loci removed | Samples removed |
| --- | --- | --- | --- |
| 10% | 250 | 14 | 1 |
| 50% | 706 | 17 | 3 |
| 90% | 790 | 69 | 13 |

### 3.3 Phylogenetic analyses and ploidy estimation

Chloroplast- and nuclear-only trees were not found to have any significant discordance; any discordance found had bootstrap values under 80 or posterior probabilities below 95, for ML and Bayesian analyses respectively. Given this, I ran analyses that concatenated both data sets for all methods and stringency levels. Despite the data being a relatively complex problem for Bayesian methodology, I was able to obtain good mixing and have analyses reach a topological convergence value of 0.01 or less after adding additional chains and lowering the temperature of the heated chains.

I found the overall topology of trees across all analyses to be fairly consistent, with one exception described further below. I found that the 10% stringency data had better mixing in the Bayesian analyses, and that results across all data sets (i.e. nuclear and chloroplast) and methods for this stringency level were more similar than those for the other two sets. Further, the support values for the 10% stringency trees were the highest. In fact, the 10% and 50% stringency sets had nearly identical topologies, with the main difference being higher support values for the former; this was also true for the coalescent methodology. The 90% stringency trees had slightly different topologies than those of the other sets, but all differences had low support values. As such, it seems that the estimations are benefiting from maximum if incomplete data, and are able to arrive at consistent, high-support topologies. For this reason, and given that Bayesian estimations have been shown to generally be more accurate (Wiens 2008), I selected the Bayesian 10% stringency tree that utilised both nuclear and chloroplast data as the ‘best tree’, and used it in subsequent analyses (Fig. 2). Select trees that exemplify trends described above can be found in SI3; all others are available upon request.

**Figure 2:**
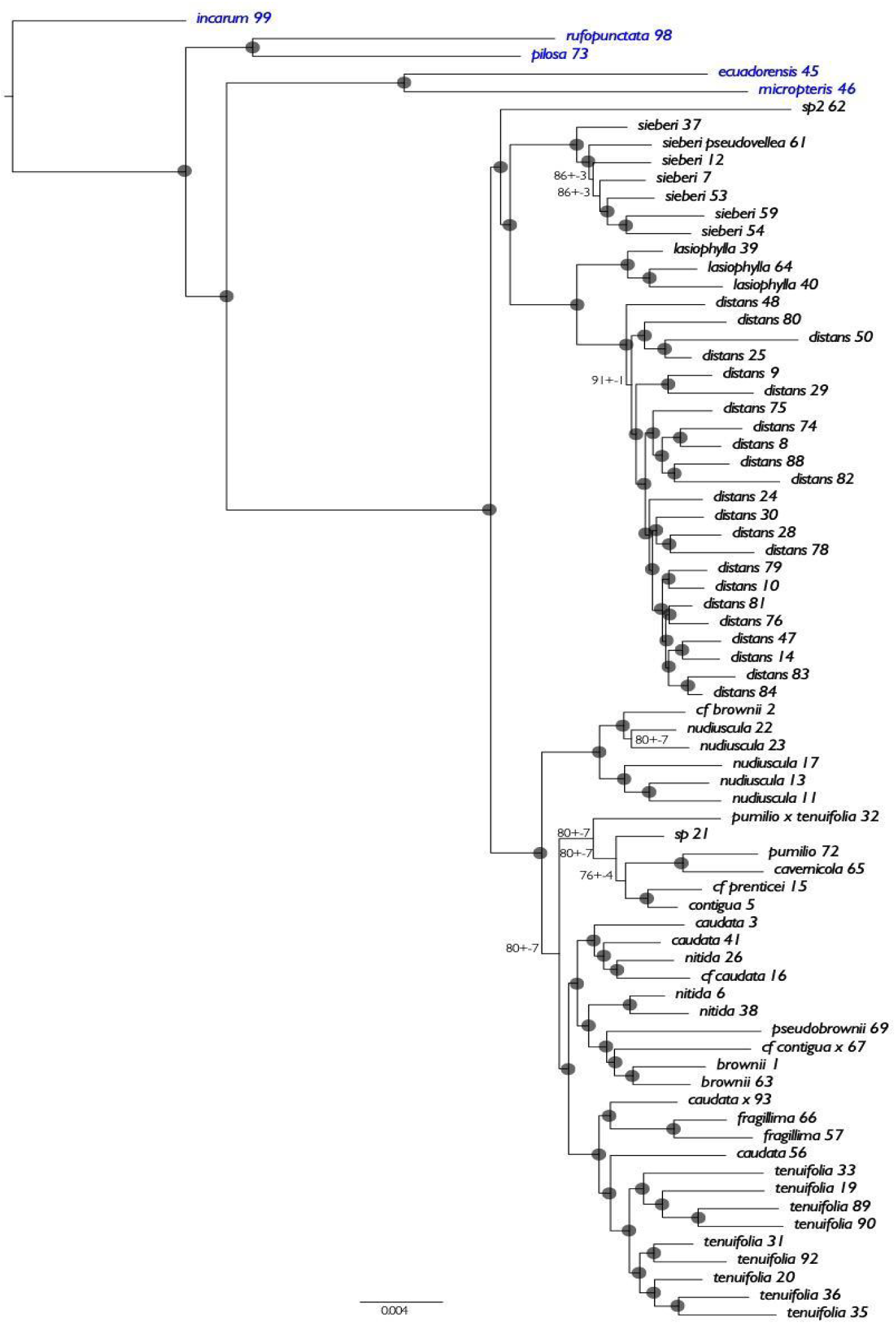
Final best tree for phylogenetic analyses, estimated through Bayesian inference and based on 10% minimum coverage with combined nuclear + chloroplast data. Grey circles signify values of 100 posterior probability ± 0 standard deviation. All values that differ from this are annotated on the tree. The species written in blue are South American, while those in black are Australasian.

Despite fairly stable topologies, I find that there is a clade—formed by *C. nitida* and *C. brownii* samples plus a handful of others—that seems to show signs of hybridisation and/or incomplete lineage sorting (this clade is the one sister to the one formed by *C. tenuifolia* and *C. fragillima* samples; Fig. 2). Although the samples always group together, this clade is unstable internally and shifts locations within its larger clade depending on the stringency scheme used, but these rearrangements always receive strong support. Because this clade belongs to the larger clade that is sister to the one *C. distans* is found in, and given that these species do not comprise a focal group, disentangling the species tree is beyond the scope of this project.

Most species formed clear monophyletic groups. Even so, there were some samples that fell outside of their species clade. I confirmed species identifications for all these samples by re-running herbarium samples through the Australian *Cheilanthes* key (Chambers and Farrant 1998). *Cheilanthes* sensu lato (which is polyphyletic in its current delimitation) has been notoriously difficult to disentangle, so misidentifications are frequent (Chambers and Farrant 1991; Quirk et al. 1983; Tryon and Tryon 1973; Ponce and Scataglini 2018). I found that I had indeed misidentified a handful of specimens, and updated their identifications on all trees. However, some species still held to their initial identification; I have kept these identifications at their original values, or added notes where I suspect possible hybrids. Disentangling Australasian *Cheilanthes* taxonomy remains an topic that warrants additional research.

I find that Australasian *Cheilanthes* are monophyletic and sister to South American *Cheilanthes*, which form the outgroup (blue species in Fig. 2). Australasian *Cheilanthes* form two sister clades. One of these includes a monophyletic *Cheilanthes tenuifolia,* along with various smaller clades formed by non-monophyletic species but whose specimens do group together across sampling schemes and analyses. One of these clades shows compelling evidence of hybridisation or incomplete lineage sorting given robust support for discordant phylogenies, as discussed above; this whole group should be subject to network analyses before the species tree can be declared with confidence. The other clade includes the target species, *C. distans*, which forms a monophyletic group sister to the monophyletic *C. lasiophylla.* Both these species form a clade sister to *C. sieberi*, which in turn are sister to what is likely a new species from Taiwan termed ‘sp2 62’ in the phylogeny. This last sample was obtained only from silica, and I was not able to see the corresponding herbarium specimen that is identified as *C. nudiuscula;* I think this determination is incorrect based on the small leaf sample used for DNA extraction and the sample’s phylogenetic placement. This and the specimen marked as ‘sp 21’—which I collected in the field and which did not readily match any species in the *Cheilanthes* key—suggest that there may be at least a couple new, undescribed species in this group.

### 3.4 Exploring reproductive mode and ploidy

I successfully estimated ploidy for 64 samples, as 10 samples yielded inconclusive results. Estimated ploidy is shown in Figure 3. All but two *C. distans* samples were found to be either triploids or tetraploids.

**Figure 3:**
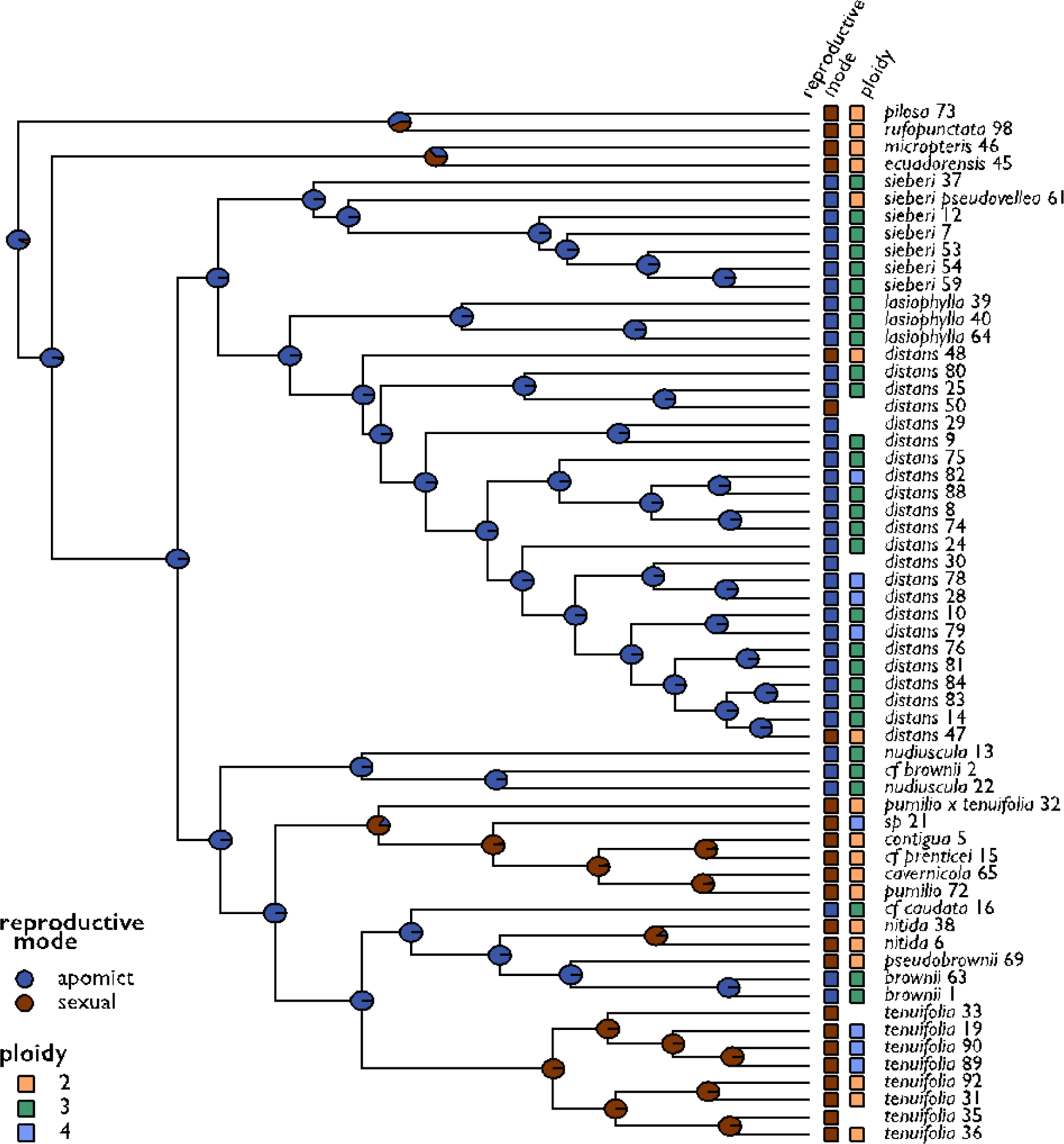
Phylogeny showing character states for reproductive mode and ploidy at the tips, and ancestral state reconstruction for reproductive mode at the nodes.

When we plot reproductive mode onto the phylogeny, we find that the presumed sexual diploid *C. distans* are in fact not monophyletic. While one sample (*distans* 48) is in fact the earliest diverging lineage in the clade, the other two (*distans* 47 and 50) are nested well within the tree (Fig. 3). I find ‘all rates different’ is the best fitting model for ancestral state reconstructions, with a log likelihood of -22.4 compared to -23.7 for ‘equal rates’. Surprisingly, even though the outgroups of the phylogeny all reproduce sexually, the ancestral state reconstruction determines the ancestor for all samples as apomictic, as well as estimating the ancestor for *C. distans* as apomictic. However, if we look at the estimated changes along the tree, a change to apomixis seems to occur specifically in Australasian *Cheilanthes* (SI2 – Fig. 2); *C. distans*, however, remains apomictic at its root. The results for the ancestral state reconstruction are certainly unusual, and their inconsistency with the stochastic character estimates (which are sampled from MCMC posterior probabilities for where on the tree character changes are occurring) may signal biases in sampling leading to erroneous estimation. Alternatively, if correct, these results may be revealing that changes in reproductive modes are more labile than previously believed. An additional round of sequencing that attempts to obtain data for the remaining presumed sexual samples, in addition to increasing sampling in apomictic *C. distans,* was performed, although ultimately not analysed; I encourage researchers to utilise these data, available at https://doi.org/10.7910/DVN/XJW1XE.

The analyses reveal all sexual specimens to be either diploids or tetraploids (Fig. 3), which is in line with prior work in this group (Tindale and Roy 2002). With regards to *C. distans*, the results do not allow us to confidently determine how many times polyploidy has arisen, especially considering that the root of the species is estimated as triploid, but we do find evidence for at least three separate shifts from triploidy to tetraploidy (SI2 – Fig. 3). The ancestral state reconstructions find the ‘all rates different’ model to have the lowest log likelihood at -54.5 compared to the ‘equal rates’ value of -56.4. Australasian *Cheilanthes* are reconstructed as triploid ancestrally, while the ancestral ploidy for the whole group remains equivocal (SI2 – Fig. 3).

The character evolution model for reproductive mode found the best fitting model to be an irreversible model of change from apomictic to sexual reproduction (Table 2), followed by the ‘all rates different’ model, then the ‘equal rates’ model, and no support for an irreversible model of changes from sexual to apomictic reproduction. Character state evolution models for ploidy found ‘equal rates’ to be the best fitting model, followed by an ordered model, a meristic model, and with an ‘all rates different’ model receiving the lowest weighted AIC score (Table 2). In the ordered model, most changes occurred from triploidy to tetraploidy (estimated transition rate = 10426.91) followed by changes from diploidy to triploidy (estimated transition rate = 27.1049).

**Table 2:** Weighted AIC values for all models tested for character evolution in reproductive mode and ploidy. Dashes signify non-applicable models.

| Character evolution model | Reproductive mode<br>(weighted AIC) | Ploidy<br>(weighted AIC) |
| --- | --- | --- |
| Equal rates | 0.117 | 0.684 |
| All rates different | 0.225 | 0.015 |
| Irreversible: sexual to apomict | 0.000 | — |
| Irreversible: apomict to sexual | 0.657 | — |
| Meristic (ordered but reversible) | — | 0.061 |
| Ordered (ordered and irreversible) | — | 0.241 |

### 3.5 Geographic patterns across the phylogeny

No phylogenetic signal beyond baseline expectation was found for spore size in *C. distans*. The estimated K statistic was 1.08 with a p-value of 0.024, meaning that specimens resemble each other as much expected under Brownian motion (the null model). The lambda statistic was estimated as 0.69 with a p-value of 0.61; while this value is not significant, it being closer to 1 means the character signal matches the phylogeny (it is not random). As such, there is as much phylogenetic signal as expected based on sample relatedness, and we cannot reject the null model that spore size is not under selection. A visual representation of the distribution of spore size across the *C. distans* phylogeny can be seen in Figure 5.

The analysis of genetic diversity by spore size groups did not find any significant difference between spore size groups. The only term found to significantly influence pi was whether a sample was tetraploid or not, with p-value=0.04 for tetraploidy and p=0.009 for the interaction between spore size group and tetraploidy (SI2 – Fig. 4).

**Figure 4:**
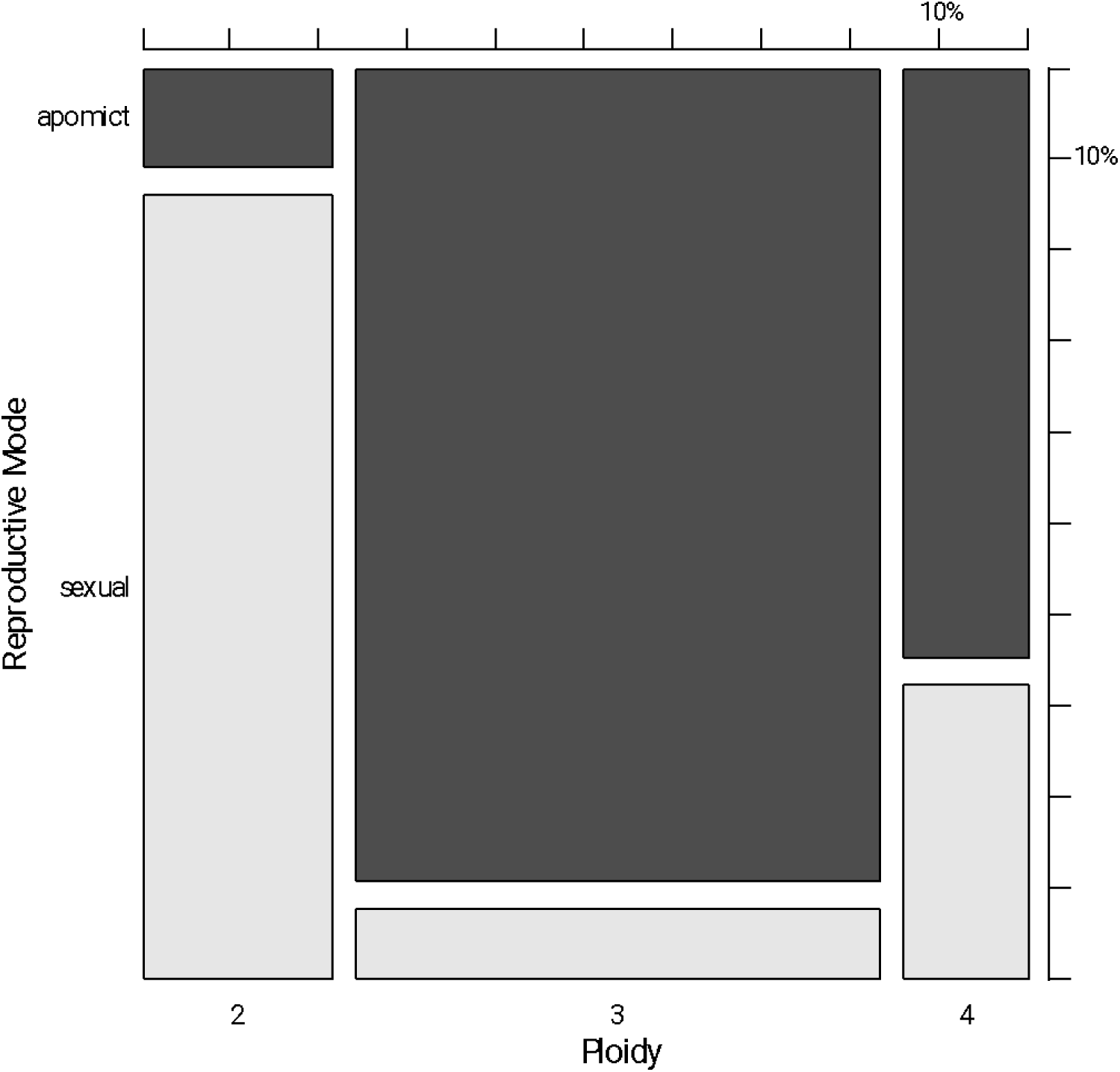
Mosaic plot for ploidy and reproductive mode (apomicts in dark grey and sexuals in light grey) for all samples. The areas of each rectangle, as well as width and height, correspond to their proportion among all samples; each tick mark on side scale bars corresponds to 10%.

**Figure 5:**
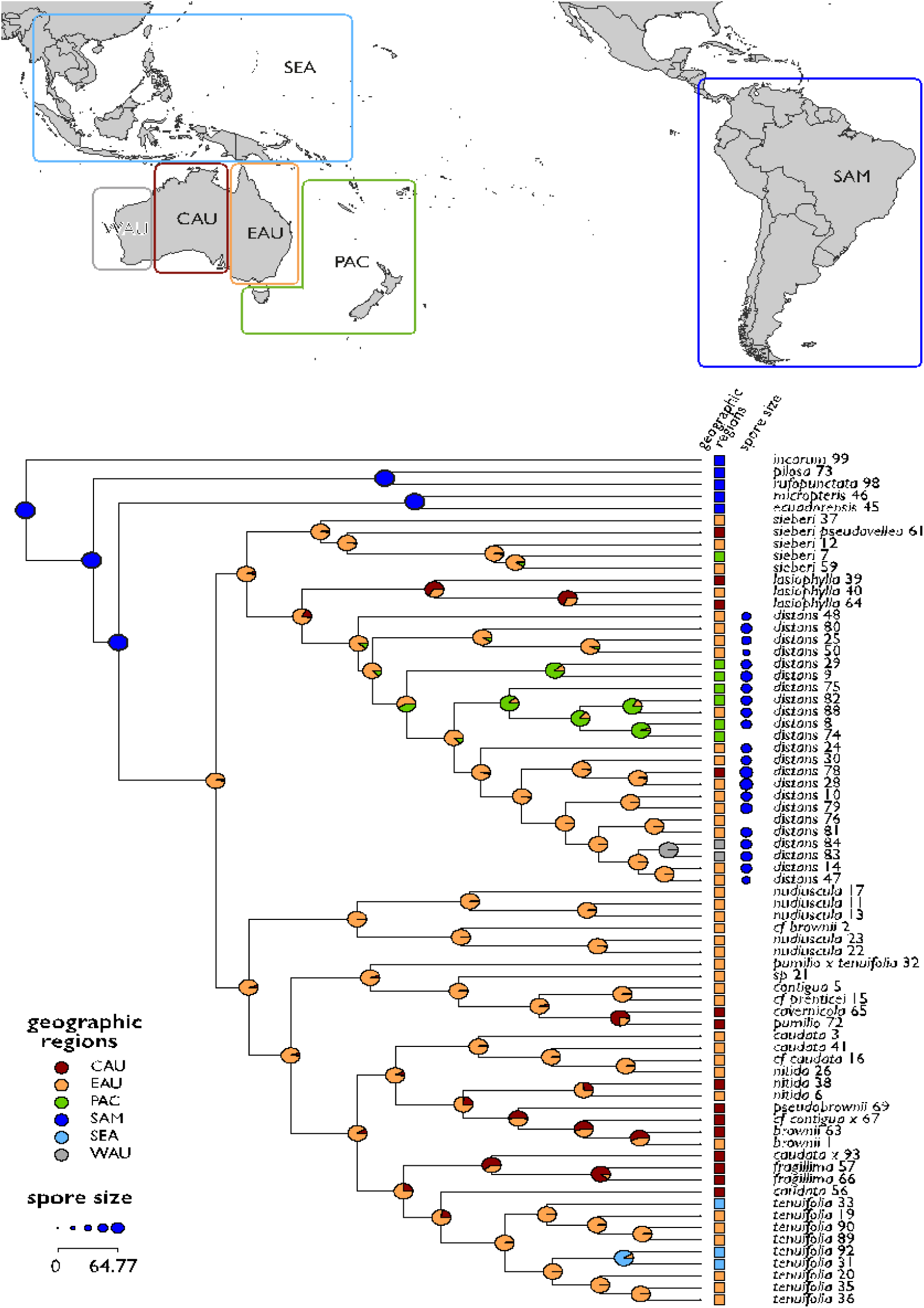
Ancestral state reconstructions for large geographic ranges. Geographic regions are defined in the map above the phylogeny and use the same colours and codes used on the tree. States at the tips are those for each specific sample. Spore sizes for *C. distans* are also displayed in blue.

Both the K statistic and lambda statistic resulted in values close to 1 for both latitude and longitude, meaning geographic distribution—even by this crude metric—seems to track the phylogeny as much as we would expect it to. In other words, the variation we observe does not reject a null hypothesis of variation being solely the product of phylogenetic relatedness. I found this result to hold when looking only at the *C. distans* samples. For the models of evolution I find that an ‘all rates different’ model that uses smaller ranges (rather than large ones) is the best fitting model for geography, with a weighted AIC score of 1 (all other models receiving a value of 0). This result is again consistent for *C. distans* only. If we look at the *C. distans* analysis, the ARD model for small regions shows that most movement occurs from Cape York Peninsula (CYK) to Eastern Australia (EAU) (estimated transition rate = 429.45), followed by Eastern Australia (EAU) to Tasmania (TAS) (estimated transition rate = 403.17), Tasmania (TAS) to New Zealand/Aotearoa (NZA) (estimated transition rate = 271.40), New Zealand/Aotearoa (NZA) to Pacific Islands (ISL) (estimated transition rate = 183.63), Cape York Peninsula (CYK) to South Australia (SAU) (estimated transition rate = 105.01), Pacific Islands (ISL) to Eastern Australia (EAU) (estimated transition rate = 84.30), and finally Eastern Australia (EAU) to Western Australia (WAU) (estimated transition rate = 35.88); all other values are barely above zero (e.g. 2.102842E-15).

The ancestral state reconstructions found ‘all rates different’ model to have the lowest log likelihood for large regions (-65.4 vs. -83.3 for ‘equal rates’), and for small regions (-96.6 vs. -133.2 for ‘equal rates’). The ancestral region for the whole group is estimated as South America for both large regions (Fig. 5) and small regions (SI2 – Fig. 5), although the probability is lower in the latter. Australasian *Cheilanthes*’ ancestral range is estimated as eastern Australia, and more specifically the Cape York peninsula. Within the whole tree analyses, *C. distans*’ ancestral state is estimated as being eastern Australia in both large and small region analyses. Stochastic character maps for both large and small regions are shown in SI2 – Figures 6 and 7.

**Figure 6:**
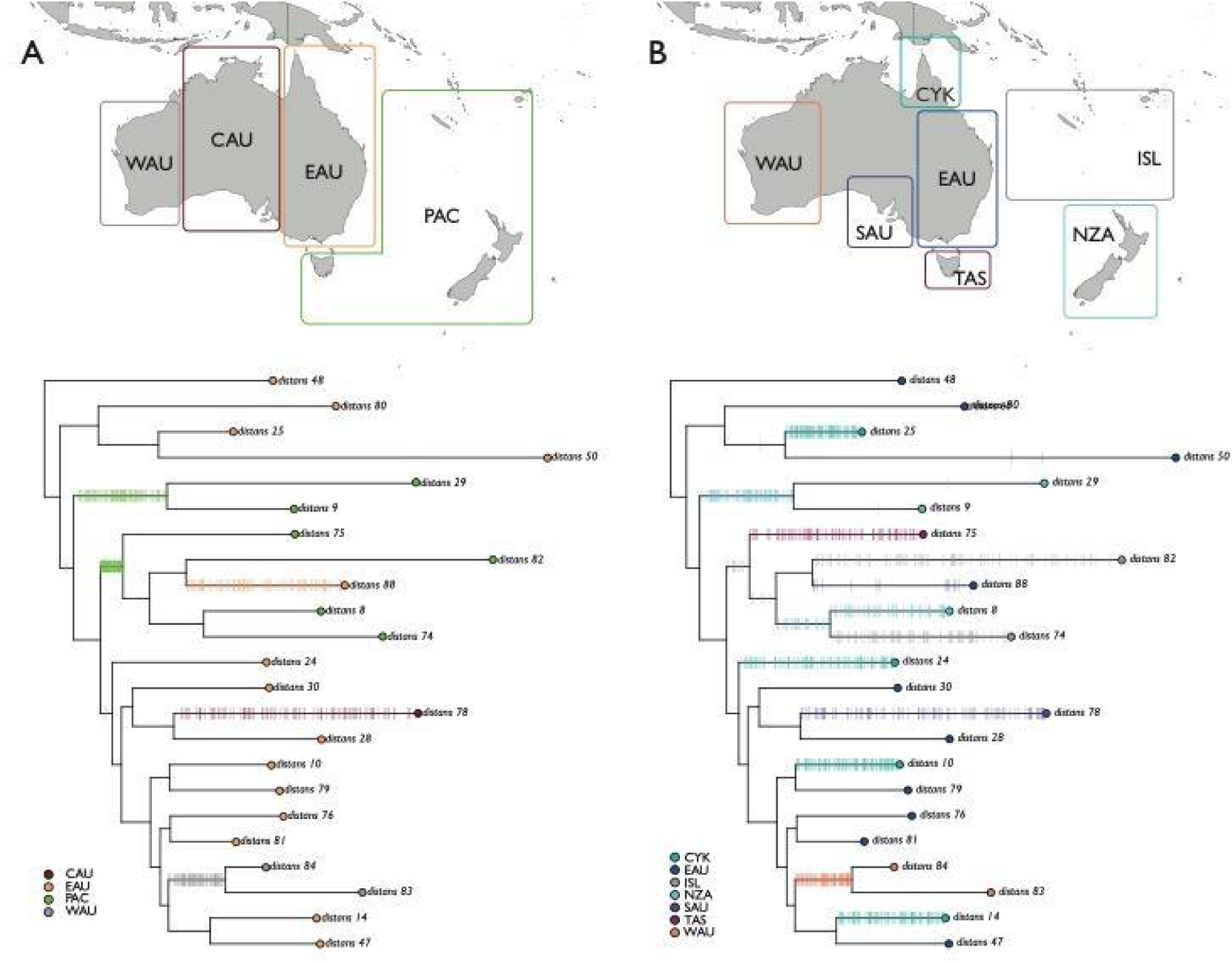
Stochastic character maps for changes in geography. The maps above the trees show the region boundaries and codes for each region, which are also used in the trees below. Each bar on the trees represents an estimated trait change based on the stochastic character estimations; a total of 100 trees sampled from the posterior probability are plotted for each of these phylogenies. A) Estimated changes when data are categorised into large regions. B) Estimated changes when data are categorised into small regions.

**Figure 7:**
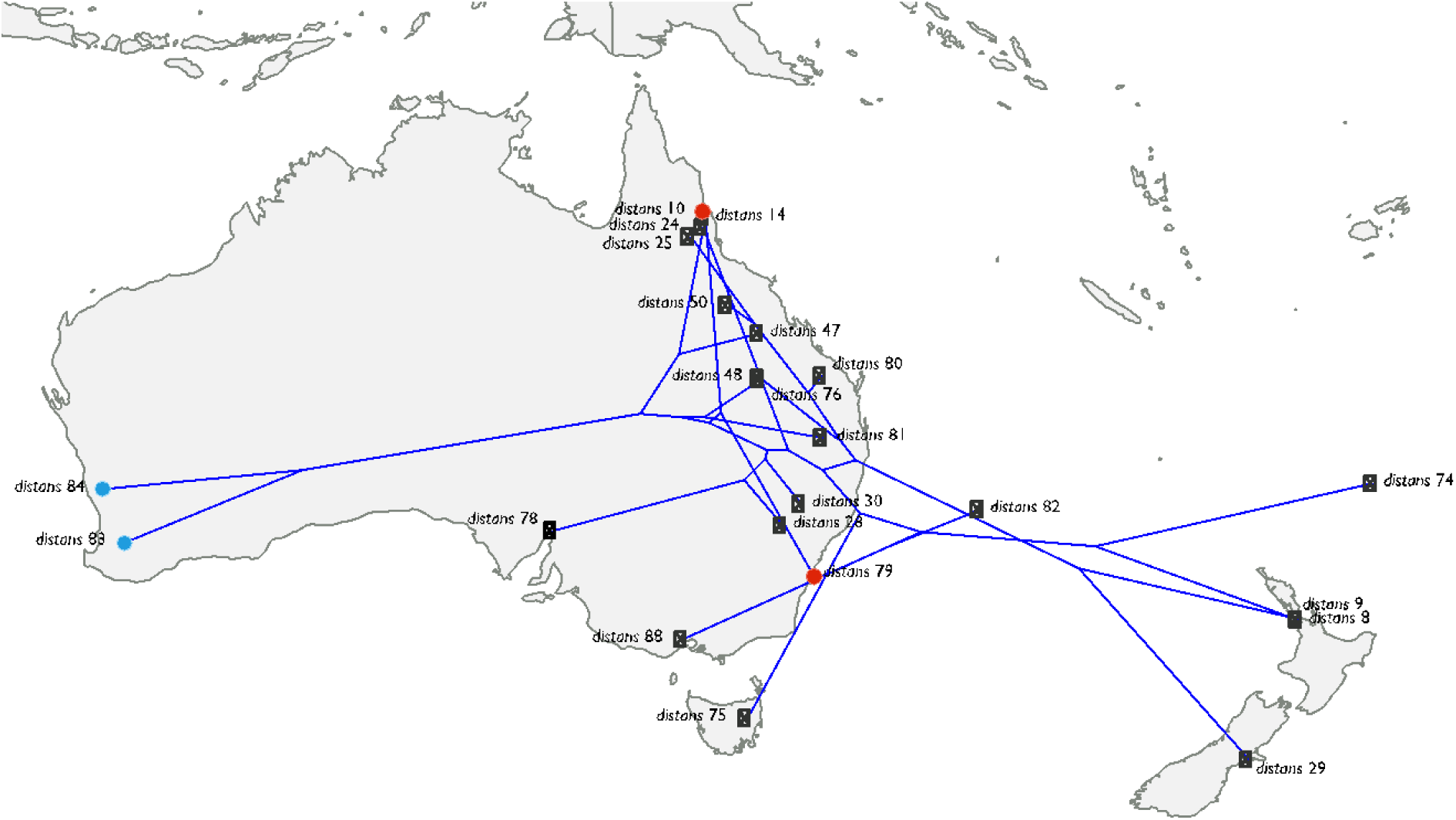
Phylogenetic tree for *Cheilanthes distans* only, plotted directly onto its geographic map. The taxa marked in blue are the sister pair with the smallest geographic distance between them, while those marked in red are the sister pair with the largest distance between them.

Ancestral state reconstructions for *C. distans-*only analyses found ‘all rates different’ model to have the lowest log likelihood for both large regions (-18.7 vs. -20.8 for ‘equal rates’) and small regions (-30.1 vs. -36.3 for ‘equal rates’). Ancestral region results for these analyses are consistent with those for the larger tree, with the exception of the ancestral state for the small region analysis recovering ancestral location as the Cape York peninsula with 75% probability, versus the near 100% probability of its ancestral range being Eastern Australia in the total tree analysis (SI2 – Fig. 5). Of particular note, however, are the patterns observed in the stochastic character maps (Fig. 6). If we look at the results for the large regions, we see that changes cluster in five branches. By contrast, changes in the small region analysis cluster in ten branches.

The plot of the *C. distans* phylogeny drawn directly onto the geographic map shows once again that lineages in this species do not appear to be restricted geographically (Fig. 7). The fact that the phylogeny has become a web that is hard to disentangle should underscore this pattern. The sister pairs with smallest and largest geographic distances have been marked onto this map.

## 4. Discussion

### 4.1 Phylogenetic analyses and ploidy estimation

Most sequence data for ferns is still limited to the chloroplast (these sample publications, among many from 2018–2022, include only plastid regions: Sessa et al. (2018); Wei et al. (2018); Chen et al. (2020); Xu et al. (2020); Fan et al. (2022)). This region has been easier to sequence and analyse because it comprises a single linkage group per specimen, reducing the complexities associated with disentangling hybrid origins or polyploid genetic diversity, both of which are especially frequent in ferns (Barrington et al. 1989; Wood et al. 2009). However, chloroplasts are maternally inherited in ferns (Gastony and Yatskievych 1992), ultimately limiting our understanding of species relationships.

Although a few nuclear markers have been developed for Pteridaceae (the most commonly used are *ApPEFP*, *gapCP*, *IBR3*, *pgiC;* Rothfels et al. (2017)), there can be considerable incongruence between the trees generated using only a handful of loci (Dyer et al. 2012; T-T. Kao, pers. comm.). One strategy to address issues of incongruence involves applying species tree methods to many loci. Yet, considering the high polyploidy and hybridity in ferns—and known polyploidy in the case of the target species of this study—most traditional high-throughput sequencing methods become inadequate, as the random and short nature of the reads generated does not allow the researcher to assess homology with confidence. This is especially true given that the only fern genomes available as references are for *Azolla* and *Salvinia* (Li et al. 2018), which are distantly related to the target *Cheilanthes*.

Target enrichment (also known as bait capture or Hyb-Seq; Weitemier et al. (2014)) has arisen as the newest technology that may offer solutions to the problems outlined above. This method preferentially sequences selected regions by using hybridization techniques, and has been successfully used to ensure acquisition of homologous data and to generate species-level phylogenies (Weitemier et al. 2014; Mandel et al. 2014; Schmickl et al. 2016; Gardner et al. 2016; Johnson et al. 2016), including its use in ferns (Wolf et al. 2017) and in polyploid species (Mandel et al. 2014; Schmickl et al. 2016). This method takes advantage of next-generation sequencing technologies that collect large amounts of data in a single sequencing run, while also targeting regions that the researcher has pre-identified as single-copy. The bait set used in this study was designed precisely to include only single-copy and low-copy regions across flagellate plants (Breinholt et al. 2021), and showed promising recovery for ferns (averaging 64.7% across all fern taxa tested).

I found high sequence recovery in my samples, averaging 93.7% recovery across nuclear loci. What is more, and in quite exciting results, recovery of chloroplast data was also high (87.4%) despite not being directly targeted by the baits; this is likely a result of the high number of chloroplasts within plant cells, but does not necessarily mean other data collection has been compromised (Schneider et al. 2021). The high recovery of loci across samples in this study is particularly exciting as it demonstrates the efficacy of a broadly-designed bait set for a variety of questions. Not only were none of the species in this study nor their close relatives included in the original bait design, but the bait design was not conceived with the specific goal of answering population-level questions. Even so, we see that enough variation in the data is recovered for a robust *C. distans* tree to be estimated. In fact, it seems that the data are comprehensive enough to reveal instances of incomplete lineage sorting and/or hybridisation, as is revealed by the *C. nitida* + *C. brownii +* allies clade; whether the data are sufficient to disentangle this complex remains to be explored. As such, sequence capture and the GoFlag 451 bait set specifically emerge as a great alternative to capture extensive data from both nuclear and plastid genomes in one single, affordable run.

Missing data in sequence alignments have been hypothesised as introducing noise and uncertainty into phylogenetic estimations, and the debate over whether fewer but more complete sequence data are better for tree estimation than more but less complete data has been ongoing for a number of years (Heath et al. 2008; Poe and Swofford 1999; Hillis et al. 2003; Pollock et al. 2002; Zwickl and Hillis 2002). My results reveal that including the most data—even if it is incomplete—results in the most robust tree estimate, as the best results are obtained with the least stringent filter of 10% minimum coverage for length, presence of loci, and presence of taxa.

In what is particularly exciting for the fern community and systematists in particular, this study presents the first phylogeny for Australasian *Cheilanthes* (Fig. 2), a group that is now confirmed as monophyletic and sister to South American *Cheilanthes*. The inclusion of all but three recognised taxa now allows the community to understand the relationship between these species, which have been particularly difficult to disentangle taxonomically (Chambers and Farrant 1991; Quirk et al. 1983; Tryon and Tryon 1973; Ponce and Scataglini 2018). This phylogeny also reveals several specimens that may represent species new to science, in addition to a previously unknown hybridisation complex involving *C. nitida* and *C. brownii,* as well as estimated ploidy data for many of the species included. It is interesting that the group that includes *C. distans, C. sieberi* and *C. lasiophylla*—all apomictic species—is stable across analyses, but that their sister clade, which includes sexual species like *C. tenuifolia,* is far less robust and shows signs of hybridisation within the group, perhaps signalling an ease for hybridisation in this group. I am hopeful that future work will allow an in-depth study of the systematics of Australasian *Cheilanthes*, including study of certain specimens of note to determine whether they represent novel species, but for now confirmation of these results remain to be explored at a later date.

### 4.2 Exploring reproductive mode and ploidy

As a result of my survey of *C. distans*, I found presumed sexual specimens in this species previously unknown to Western science (see Sosa 2024). Workers have found that apomictic polyploid fern lineages are most often derived from diploid sexual progenitors (Beck et al. 2010; Sigel et al. 2011; Dyer et al. 2012), especially since the reversal from apomixis—for triploids in particular—to sexually reproducing diploids is at the very least theoretically difficult, having to accomplish several unlikely steps than would a change from sexual to apomictic reproduction (Grusz 2016; Grusz et al. 2021). I wished to explore whether the presumed sexual specimens I found were in fact the parental lineage to apomictic *C. distans.* Further, it is of interest whether polyploid lineages have arisen more than once as multiple origins transfer a greater proportion of genetic variation, especially so into apomictic lineages, potentially enhancing their evolutionary success. In fact, several studies have shown that it is common for polyploids to arise more than once (Werth et al. 1985; Soltis et al. 2004; Cifuentes et al. 2010; Beck et al. 2012; Sigel et al. 2014; Wickell et al. 2017).

Surprisingly, we find that the sexual diploid *C. distans* specimens are not monophyletic, and thus do not seem to form a single parental lineage to apomictic members of this species. It is possible that there are sexual progenitors to all apomictic lineages, as one sexual sample is the earliest diverging lineage of the entire group (i.e. *distans* 48). Nevertheless, the other two samples are nested within the apomictic samples. This does not necessarily mean that these sexual lineages have arisen *from* apomictic lineages, since biases in data could lead to these results. In fact, Cook and Crisp specifically outline how incorrect results precisely like these may be obtained in cases where frequent dispersals happen asymmetrically from a narrow parental range, resulting in analyses that reconstruct the ancestral range as a terminal node in the tree (Cook and Crisp 2005). Further evidence for biassed results arises from the fact that sexual specimens only occur in a relatively narrow range, and the apomictic lineages with which they are closest phylogenetically are not geographically close to this range (the apomictic samples being located 460 to 3000km away). Although one could imagine a scenario where a diploid lineage arises within a particularly populous apomictic population, both the geographic distances and known tree reconstruction biases give us reasons to doubt that this is what the present data show.

However, if the results are taken at face value, they point to the possibility of reversal from apomictic triploid lineages to diploid sexual lineages. It is possible that the environmental conditions faced by a xeric-adapted *C. distans* may be affecting meiosis and reproduction to such an extent that reversals become more likely, as we know that harsh conditions affect sporogenesis (De Storme and Geelen 2014 and references therein for studies on pollen; Cox and Hickey 1984; Arosa et al. 2009). Although to my knowledge this has not been reported before in ferns, we should not discount this possibility outright. In fact, the results from the ancestral state reconstruction support this hypothesis, as they suggest that the ancestral state for the group and for *C. distans* is apomixis, and phylogenetic signal points to reproduction as being a highly conserved trait. Further, these findings are supported by the fact that the best fitting character evolution model is one of irreversible change from apomixis to sexual reproduction. There is, however, some discordance when we look at the stochastic character maps (generated from sampling the posterior probability that estimates along which branches changes occur) given that reproductive mode change in this analysis is estimated to occur at the base of Australasian *Cheilanthes* rather than being the ancestral state for the whole group (SI2 – Fig. 2); this discordance may be pointing to bias in the tree that perhaps drives the results we see in our estimations. Regardless, the exact dynamics leading to changes in reproductive mode in *C. distans* deserve more detailed study.

Unfortunately, the particular structure of the current tree does not allow us to obtain an estimate on the number of origins for polyploid lineages for *C. distans*, with the exception of origins of tetraploid lineages, for which there have been at least three origins. The reconstruction of the ancestral state of this species as triploid is surprising, but is in line with the reproductive mode results outlined above. That the best fitting character evolution model is ‘equal rates’—meaning that all changes from one ploidy to another are equally likely—is also somewhat unusual given theoretical expectations of ploidal reductions being harder to achieve than ploidal additions. Analyses of chromosomal arrangements and/or total genomic content should provide insights into whether these results are due to biases in sampling or are in fact revealing reversal events.

Regardless of the uncertainties outlined above, it is exciting that the data do robustly point to support an association between ploidy and reproductive mode, as has been reported by previous workers (Tindale and Roy 2002; Manton 1950; Walker 1962; Windham and Yatskievych 2003). I find that sexual specimens are either diploids or tetraploids (Fig. 3), and that sexual reproduction itself is associated especially with diploidy and—more weakly—with tetraploidy (Fig. 5). Apomixis is mostly associated with triploidy, a result that has also been found in other ferns (Grusz 2016; Grusz et al. 2021). Given the small difference found between tetraploidy and reproductive mode, as well as considering previous equivocal findings, exploring the association between tetraploidy and reproductive mode remains an area that deserves further exploration.

### 4.3 Geographic patterns across the phylogeny

It is unexpected that fern population genetics have not received much attention, and that certain geographic regions in particular have almost no data (Pelosi and Sessa 2021): for Australia and New Zealand/Aotearoa, only two out of 508 taxa in this area have been studied (Shepherd et al. 2007; Keiper and McConchie 2000). Apomictic ferns have also been particularly understudied, with only a handful of publications that explore their population genetics (Haufler 1985; Watano and Iwatsuki 1988; Lin et al. 1995) and only two that look at DNA sequence data rather than relying on electrophoretic analyses (Wickell et al. 2017; Ootsuki et al. 2011). Yet population-level data are necessary to answer a variety of questions, including understanding geographic distributions and dispersal dynamics in a group. Ferns in particular have been hypothesised as being able to disperse over large distances, with their abundance on isolated islands often cited as proof (Tryon 1970; Smith 1972; Peck et al. 1990). However there have been no studies—given the dearth of appropriate data—that analyse the population structure of widely-distributed species with varying reproductive modes and ploidies, assess their geographic distributions, and test whether they have attained their ranges through many small steps or a few larger jumps.

The most salient result of my analyses is perhaps the finding that most dispersal in *C. distans* is occurring over shorter distances rather than always occurring over long jumps. This is evidenced by the fact that we see double the number of transitions in geography when regions are defined narrowly vs. when I use larger regions for the analyses (Fig.6). In other words, there are half the number of dispersal events between large regions—where dispersal must also most often happen over longer distances in order to ‘escape’ the larger region—than there are dispersal events between smaller regions. Shorter distance movement being more frequent is also supported by the fact that the best fitting evolution models are those that code the data into smaller regions, even as these models have been weighted down due to their higher number of parameters. Thus we also see character evolution models recover dispersal as occurring over smaller distances rather than longer ones. Although we might expect shorter distance dispersal is more frequent than long distance dispersal, this is the first study to provide concrete data demonstrating this dynamic in ferns, and over a large area in a widely distributed species. A previous study had found significant empty habitat for a specialised fern (Wild and Gagnon 2005); these findings support the possibility that fern spores are not as dispersible as has been posited by the fern community in the past.

Further, there has been long standing interest in understanding the role of winds in the dispersal of organisms (Sanmartín et al. 2007; Muñoz et al. 2004). Excitingly, my analyses also reveal that most dispersal seems to be occurring from Cape York Peninsula to Eastern Australia, from Eastern Australia to Tasmania, from Tasmania to New Zealand/Aotearoa, and from New Zealand/Aotearoa to other islands in the Pacific. These movements would seem to track trade winds, especially those in the summer (Edward and Bart 1996) when most spores are being produced. However, plenty of spores remain on leaves past the growing season (Sosa, pers. obvs.), and these may also be carried in the directions predicted by the analyses during subtropical ridges in the winter (BoM 2022).

For widely-distributed species such as *C. distans*, we can also ask whether their large ranges have resulted from a subset of particularly dispersible lineages, or whether all lineages are more or less equally successful at dispersal. In the case of *C. distans*, I do not find evidence of certain lineages clustering geographically—or, put otherwise, being particularly poor dispersers—nor do I find lineages that are over-dispersed in space—put otherwise, being particularly *good* dispersers. The plot of the *C. distans* phylogeny directly onto geographic space (Fig. 7) confirms the finding that dispersal among the lineages of this species does not seem to be limited geographically. This also means that *C. distans*’ range is not a composite of smaller ranges of isolated lineages, as has been found in other xeric-adapted ferns (Beck et al. 2012; Wickell et al. 2017), but rather a single large range. This might then be a case supporting the General Purpose hypothesis, where polyploidy has provided the genetic variation necessary to survive and reproduce in a variety of environments (Lynch 1984); further investigation into niche variation across *C. distans’* range is necessary to confirm this hypothesis.

Another interesting question we can ask is whether spore size is correlated by lineage, especially given my findings that spore size in *C. distans* may be influencing both range size and germination (described in Sosa 2024). If spore size were phylogenetically structured in such a way that spore sizes group by lineage, and considering my earlier findings that medium spores attain the largest ranges, it would mean that certain lineages are likely better dispersers than others. However, I do not find this to be the case, and instead find that spore size variation does not differ beyond what we would expect as a result of phylogenetic relatedness. My results stand in some contrast with findings by previous workers, who noted what they thought was greater than expected randomness in spore formation, since they observed that spore mother cells underwent meiosis more or less independently of each other and that each sporangium within a single plant could result in different types and numbers of spores (Ekrt and Koutecký 2016; Mehra and Singh 1957; Bell 1960; Manton 1950). Understanding in more detail how conserved spore size is as a character remains an area requiring further exploration.

If a certain spore size group is in fact dispersing into larger ranges, we might possibly expect this group to have higher genetic diversity. This could arise either by these lineages gaining additional genetic variation through low levels of sexual reproduction (since apomicts are known to produce viable sperm (Walker 1962, 1985)) or, if we assume only asexual reproduction, then if most of the variation in spore size arose from frequent generation of novel lineages, where some of these lineages are more successful at dispersing due to their spore size. However, I find that genetic diversity is more or less the same for all spore size groups, with any significant changes being due solely to ploidy increase (especially from triploidy to tetraploidy). This observation, coupled with the lack of phylogenetic signal for spore size, once again underscore a lack of conservation of spore size within a lineage.

The present data also allow us to gain a better understanding of the biogeography of Australasian *Cheilanthes,* which had previously not been possible as no phylogeny existed for the group. I find the whole group has arisen from and is nested within South American relatives. Dispersal from South America to this region is estimated as having happened to eastern Australia, possibly as specifically as the Cape York peninsula area. This area is also the one in which *C. distans* itself originates. The high biodiversity of the Cape York peninsula is already well known (Crisp et al. 2001). If this region is in fact the centre of origin for the group and for several of its species, it could support previous hypotheses of Cape York being a refugium that provides favourable conditions over extended periods of time and thus allows for increased diversification (VanDerWal et al. 2009; for examples demonstrating higher diversification within refugia see Murphy et al. 2015; and Condamine et al. 2017).

## 5. Conclusion

The importance of population-level data are revealed in this study, especially for particularly understudied groups and geographic regions. I was able to obtain sequence data across hundreds of nuclear and dozens of plastid loci, and with these build the first ever phylogeny for Australasian *Cheilanthes*. This phylogeny included population-level sampling for *C. distans*, and with this tree I was able explore trait evolution and geographic dispersal. I estimate the ancestral region for *Cheilanthes*, and find evidence for Cape York potentially playing an important role as a refugium and place of diversification within this genus. When looking at dynamics within *C. distans*, I find that sexual specimens of this species do not appear to be monophyletic, nor the parental lineage to apomict specimens, although whether these results are due to bias or are in fact showing the first evidence in ferns for reversal from apomixis to sexual reproduction remains to be explored. I also find evidence for ploidy and reproductive mode being tightly linked. Excitingly, I also find compelling evidence that most dispersal in *C. distans* is occurring over shorter rather than longer distances, underscoring how short movement can nevertheless allow for establishment of large ranges. Further, dispersal does not seem to be restricted by lineage, and seems to be occurring more or less evenly across the tree. What is more, I find that dispersal shows evidence of being asymmetric and tracking trade winds, adding concrete data to hypotheses that have mostly been explored theoretically up until now.

## Acknowledgements

First and foremost, apologies and amends are owed to the Traditional Custodians and Indigenous Nations from whose lands samples in this study were taken without consent nor compensation. While I have done my best to be respectful of their non-human relatives (i.e. the ferns studied herein), I have certainly fallen short.

This research would not have been possible without generous funding from the following institutions and organisations: the American Society of Plant Taxonomists, the Biology Department at Duke University, the Botanical Society of America, the International Association for Plant Taxonomy, the National Geographic Society, Sigma Xi, and the Society of Systematic Biologists. Fieldwork was possible thanks to the National Geographic Society’s Young Explorers Grant. I am additionally indebted to all the herbaria who graciously lent me their specimens: State Herbarium of South Australia (AD), Auckland War Memorial Museum (AK), Natural History Museum – UK (BM), Queensland Herbarium (BRI), Australian National Herbarium (CANB), Manaaki Whenua – Landcare Research (CHR), Australian Tropical Herbarium (CNS), Department of Environment, Parks and Water Security (DNA), James Cook University (JCT), Tasmanian Museum and Art Gallery (HO), Royal Botanic Gardens Victoria (MEL), Missouri Botanical Garden (MO), University of New England (NE), Royal Botanic Gardens and Domain Trust (NSW), Western Australian Herbarium (PERTH), National Tropical Botanical Garden (PTBG), Smithsonian Institution (US), and Taiwan Forestry Research Institute (TAIF).

I am incredibly grateful to Mark Rausher for welcoming me into his lab and for his steadfast support. His insightful thoughts, along with those of Anne Yoder, Louise Roth, and Paul Manos, significantly improved this work. I owe much gratitude to innumerable colleagues and friends for asking astute questions and making keen observations that have much improved my science; special thanks are due to Ray Allen, Lauren Carley, Jen Coughlan, Tzu-Tong Kao, Devin McCarthy, and Kathryn Picard. An immense debt is owed to Layne Huiet for her assistance with the loans involved in this study, to Ashley Field for his assistance with fieldwork, to Gordon Burleigh for his support with sequencing, to George Tiley and Matt Johnson for their help with questions regarding phylogenetics and bioinformatics, and to Julia Notar for her help in reviewing and formatting this paper for publication.

Some of the ideas in this study were first developed under tutelage of Kathleen Pryer and Michael Windham.

