## Supplemental Information 1 for "Short steps can take you far: Phylogenetic analysis of Australasian *Cheilanthes distans* reveals frequent shorter-range dispersal"

**Table SI1.1: Samples selected for phylogeny generation. Notes have been made on DNA source material, with 'KS silica' denoting samples I collected in the field. Whether extractions and/or sequencing were successful has been recorded for each sample.**

| Seq. ID | *Cheilanthes* species | Herbarium ID; Collector & No. | Material source | Extraction successful? | Sequencing successful? |
|---|---|---|---|---|---|
| 58 | *adiantoides* | NSW 363288; RJ Chinnock 6703 | silica | yes | no |
| 68 | *adiantoides* | MEL 1615273; RJ Chinnock 6703 | herbarium | yes | yes |
| 1 | *brownii* | DUKE; K Sosa 9 | KS silica | yes | yes |
| 42 | *brownii* | MO 6125469; van d. Werff 22406 | herbarium | yes | no |
| 63 | *brownii* | MO 6125469; van d. Werff 22406 | silica | yes | yes |
| 3 | *caudata* | DUKE; K Sosa 45 | KS silica | yes | yes |
| 41 | *caudata* | NSW 668596; J Kemp TH444 | herbarium | yes | yes |
| 56 | *caudata* | NSW 238573; GJ Leach 2751 | silica | yes | yes |
| 93 | *caudata x* | CANB 349227.1; DL Jones 1521 | herbarium | yes | yes |
| 65 | *cavernicola* | MEL 2032290; NG Walsh 4451 | silica | yes | yes |
| 16 | *cf .caudata* | DUKE; K Sosa 33 | KS silica | yes | yes |
| 2 | *cf. brownii* | DUKE; K Sosa 11 | KS silica | yes | yes |
| 67 | *cf. contigua x* | MEL 2036136; MF Duretto 1005 | herbarium | yes | yes |
| 15 | *cf. prenticei* | DUKE; K Sosa 32 | KS silica | yes | yes |
| 34 | *cf. tenuifolia* | BRI AQ0679002; PI Forster PIF24782 | herbarium | no | no |
| 95 | *cf. tenuifolia* | MEL 1595692A; BD Duncan s.n. | herbarium | no | no |
| 96 | *cf. tenuifolia* | MEL 0564690A; TB Muir 6101 | herbarium | no | no |

| | | | | | |
|---|---|---|---|---|---|
| 4 | *contigua* | DUKE;<br>K Sosa 31 | KS silica | yes | no |
| 5 | *contigua* | DUKE;<br>K Sosa 34 | KS silica | yes | yes |
| 8 | *distans* | DUKE;<br>K Sosa 1 | KS silica | yes | yes |
| 9 | *distans* | DUKE;<br>K Sosa 5 | KS silica | yes | yes |
| 10 | *distans* | BRI AQ1006867;<br>K Sosa 10 | KS silica | yes | yes |
| 14 | *distans* | BRI AQ1006877;<br>K Sosa 20 | KS silica | yes | yes |
| 24 | *distans* | BRI AQ1006889;<br>K Sosa 55 | KS silica | yes | yes |
| 25 | *distans* | BRI AQ1006892;<br>K Sosa 58 | KS silica | yes | yes |
| 28 | *distans* | NSW 851335;<br>BM Wiecek 1648 | herbarium | yes | yes |
| 29 | *distans* | CHR 383122;<br>G Brownlie 1103 | herbarium | yes | yes |
| 30 | *distans* | CANB 871782.1;<br>I Crawford 8708 | herbarium | yes | yes |
| 47 | *distans* | BRI AQ0743349;<br>RJ Fensham 5341 | herbarium | yes | yes |
| 48 | *distans* | BRI AQ0628082;<br>D Halford Q2100 | herbarium | yes | yes |
| 49 | *distans* | BRI AQ0415458;<br>RW Johnson<br>2152A | herbarium | no | no |
| 50 | *distans* | BRI AQ0767605;<br>G Sankowsky<br>2392 | herbarium | yes | yes |
| 51 | *distans* | BRI AQ0565430;<br>RJ Fensham 317 | herbarium | no | no |
| 52 | *distans* | NSW 192038;<br>unknown | herbarium | no | no |
| 70 | *distans* | BRI AQ0224313;<br>RW Johnson 1261 | herbarium | no | no |
| 74 | *distans* | AK325810;<br>PJ de Lange s.n. | herbarium | yes | yes |
| 75 | *distans* | HO 316224;<br>AM Gray 788 | herbarium | yes | yes |
| 76 | *distans* | NE 105628;<br>STS Ling 18 | herbarium | yes | yes |
| 77 | *distans* | CANB 45260.1;<br>R Pullen 661 | herbarium | yes | no |

| | | | | | |
|---|---|---|---|---|---|
| 78 | *distans* | AD 99505280; DE Mufret 49-1088 | herbarium | yes | yes |
| 79 | *distans* | CANB 631929.1; DL Jones 17762 | herbarium | yes | yes |
| 80 | *distans* | BRI AQ0836645; PI Forster PIF39826 | herbarium | yes | yes |
| 81 | *distans* | BRI AQ0776801; AR Bean 20341 | herbarium | yes | yes |
| 82 | *distans* | BRI AQ0643352; JM Le Cussan 981 | herbarium | yes | yes |
| 83 | *distans* | PERTH 6593321; G Warren 721 | herbarium | yes | yes |
| 84 | *distans* | PERTH 6070515; M Henson MJH 39 | herbarium | yes | yes |
| 85 | *distans* | CANB 176136.1; RDHoogland 11309 | herbarium | no | no |
| 86 | *distans* | NSW 191947; AN Rodd 1757 | herbarium | no | no |
| 87 | *distans* | NSW 192043; HS McKee 2424 | herbarium | yes | no |
| 88 | *distans* | MEL 2021001A; V Stajsic 872 | herbarium | yes | yes |
| 45 | *ecuadorensis* | Madsen & Sanches 7940 | herbarium | yes | yes |
| 57 | *fragillima* | NSW 241335; CR Dunlop 8625 | silica | yes | yes |
| 66 | *fragillima* | MEL 260166; CR Dunlop 10039 | herbarium | yes | yes |
| 99 | *incarum* | UC 1781972; M Lehnert 340 | herbarium | yes | yes |
| 39 | *lasiophylla* | NSW 222447; MJ Barritt 30 | herbarium | yes | yes |
| 40 | *lasiophylla* | NSW 26609; W Greuter 20743 | herbarium | yes | yes |
| 64 | *lasiophylla* | MEL 2201097; Latz 17080 | herbarium | yes | yes |
| 46 | *micropteris* | H. Huaylla 69 | herbarium | yes | yes |
| 6 | *nitida* | DUKE; K Sosa 8 | KS silica | yes | yes |
| 26 | *nitida* | DUKE; K Sosa 38 | herbarium | yes | yes |

| | | | | | |
|---|---|---|---|---|---|
| 38 | *nitida* | NSW 232678; CR Dunlop 8612 | herbarium | yes | yes |
| 11 | *nudiuscula* | DUKE; K Sosa 12 | KS silica | yes | yes |
| 13 | *nudiuscula* | DUKE; K Sosa 17 | KS silica | yes | yes |
| 17 | *nudiuscula* | DUKE; K Sosa 40 | KS silica | yes | yes |
| 22 | *nudiuscula* | DUKE; K Sosa 59 | herbarium | yes | yes |
| 23 | *nudiuscula* | DUKE; K Sosa 56 | herbarium | yes | yes |
| 91 | *nudiuscula* | MEL 2035733; MF Duretto 897 | herbarium | yes | yes |
| 100 | *obducta* | DUKE 400686; FM Zardini 50137 | herbarium | yes | no |
| 73 | *pilosa* | DUKE 401862; CF Rothfels 3980 | silica | yes | yes |
| 43 | *praetermissa* | NSW 624046; RW Johnson 4646 | herbarium | no | no |
| 44 | *praetermissa* | MO 3261415; DJ Dixon 1373 | herbarium | yes | no |
| 60 | *prenticei* | NSW 218098; BL Rice 2412 | silica | no | no |
| 69 | *pseudobrownii* | NSW 586734; A Fraser 284 | herbarium | yes | yes |
| 18 | *pumilio* | DUKE; K Sosa 43 | KS silica | yes | no |
| 72 | *pumilio* | MEL 2036133; MF Duretto 1038 | herbarium | yes | yes |
| 55 | *pumilio* | NSW 217703; J R-Smith 8064 | silica | no | no |
| 32 | *pumilio x tenuifolia* | BRI AQ0468957; PD Bostock 642 | herbarium | yes | yes |
| 98 | *rufopunctata* | MO 6124754; L Valenzuela 8350 | herbarium | yes | yes |
| 71 | *shirleyana* | BRI AQ0224383; Gray s.n. | herbarium | no | no |
| 7 | *sieberi* | DUKE; K Sosa 4 | KS silica | yes | yes |
| 12 | *sieberi* | DUKE; K Sosa 13 | KS silica | yes | yes |
| 37 | *sieberi* | BRI AQ1006891; K Sosa 57 | herbarium | yes | yes |
| 53 | *sieberi* | DUKE; Nagalingum 20 | silica | yes | yes |

| | | | | | |
|---|---|---|---|---|---|
| 54 | *sieberi* | DUKE; Nagalingum 22 | silica | yes | yes |
| 59 | *sieberi* | NSW 257216; S Bell YUT92/287 | silica | yes | yes |
| 94 | *sieberi* | MEL 1595682A; BD Duncan s.n. | herbarium | yes | yes |
| 61 | *sieberi subps. pseudovellea* | NSW 586735; A Fraser 281 | silica | yes | yes |
| 21 | *sp* | DUKE; K Sosa 54 | KS silica | yes | yes |
| 62 | *sp2* | TAIF; LY Kuo 211 | silica | yes | yes |
| 19 | *tenuifolia* | DUKE; K Sosa 6 | KS silica | yes | yes |
| 20 | *tenuifolia* | DUKE; K Sosa 50 | KS silica | yes | yes |
| 27 | *tenuifolia* | BRI AQ1008110; K Sosa 27 | KS silica | yes | no |
| 31 | *tenuifolia* | DUKE; Grusz 1 | herbarium | yes | yes |
| 33 | *tenuifolia* | BRI AQ0507571; M Card CAP3 | herbarium | yes | yes |
| 35 | *tenuifolia* | BRI AQ1006876; K Sosa 19 | KS silica | yes | yes |
| 36 | *tenuifolia* | BRI AQ1008123; K Sosa 51 | KS silica | yes | yes |
| 89 | *tenuifolia* | BRI AQ0836424; PI Forster PIF40070 | herbarium | yes | yes |
| 90 | *tenuifolia* | BRI AQ0836288; PI Forster PIF39985 | herbarium | yes | yes |
| 92 | *tenuifolia* | CANB 215822.1; K Paijmans 524 | herbarium | yes | yes |
