## Supplemental Information 2 for "Short steps can take you far: Phylogenetic analysis of Australasian *Cheilanthes distans* reveals frequent shorter-range dispersal"

This appendix contains some additional figures and tables, with the exception of phylogenetic inference results, which are included in SI3. See the main text for context.

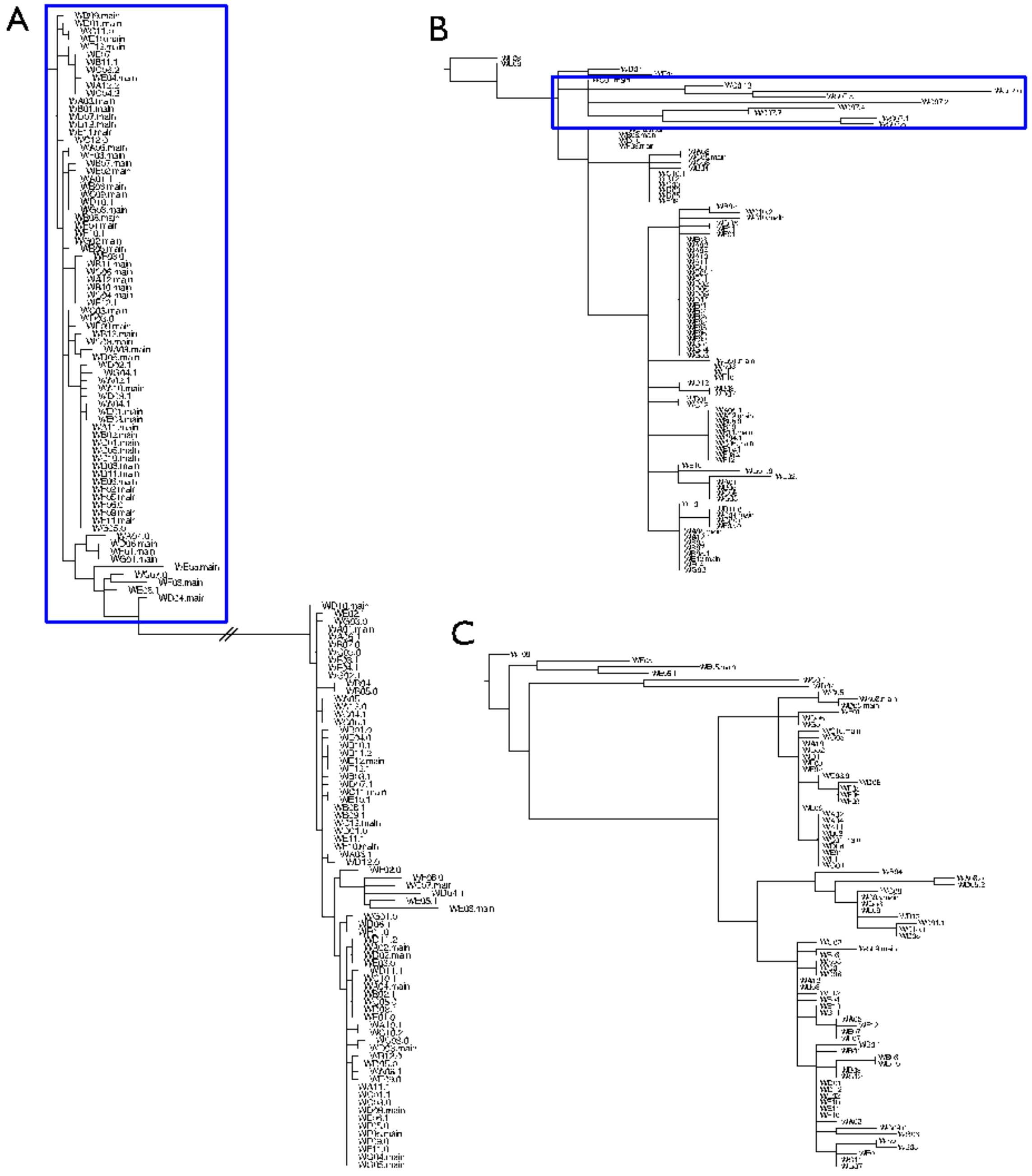


**Figure SI2.1: Types of paralog trees observed. A) Clear structure shows gene duplication. Sequences selected in these trees were those belonging to the clade with most 'main' paralogs, as those show the highest quality; for this example, this clade is marked in blue. B) Paralogs cluster into a monophyletic group, marked in this tree in blue. C) No clear structure observed with regards to paralog location on the phylogeny.**

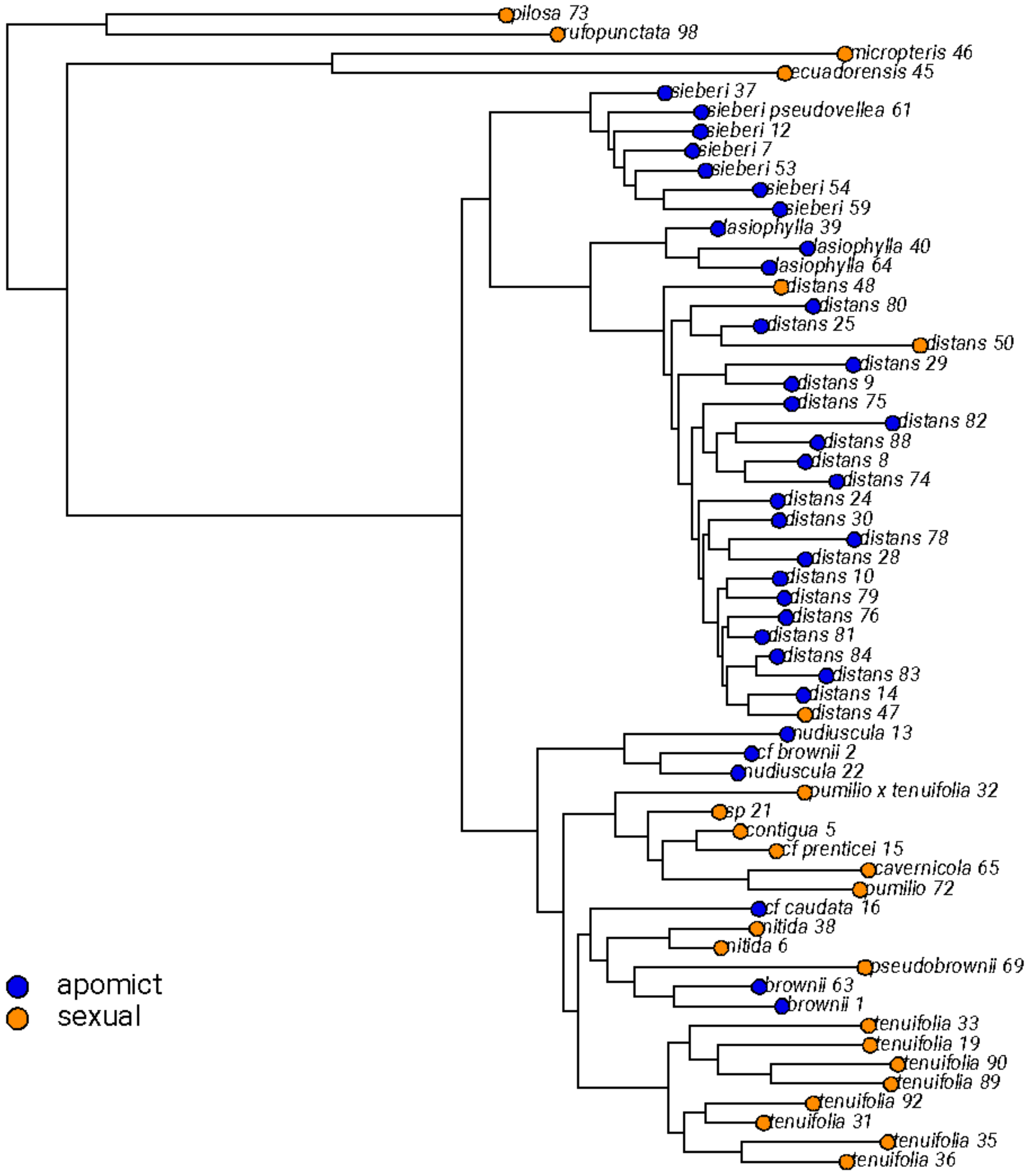


**Figure SI2.2: Stochastic character changes for reproductive mode, based on 100 sampled trees. Each bar represents an estimated change; a total of 100 trees were sampled from the posterior and are plotted onto this phylogeny.**

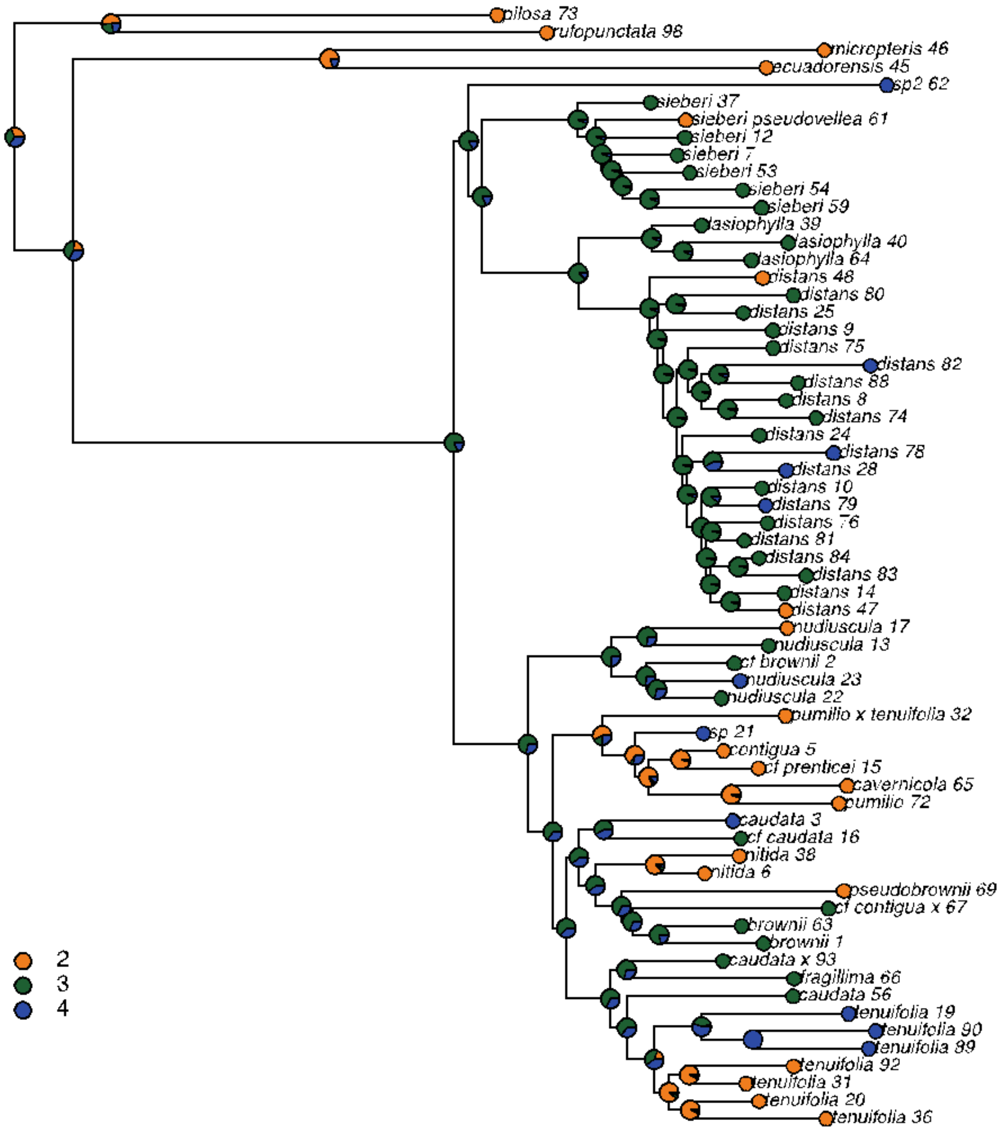


**Figure SI2.3: Ancestral state reconstructions for ploidy.**

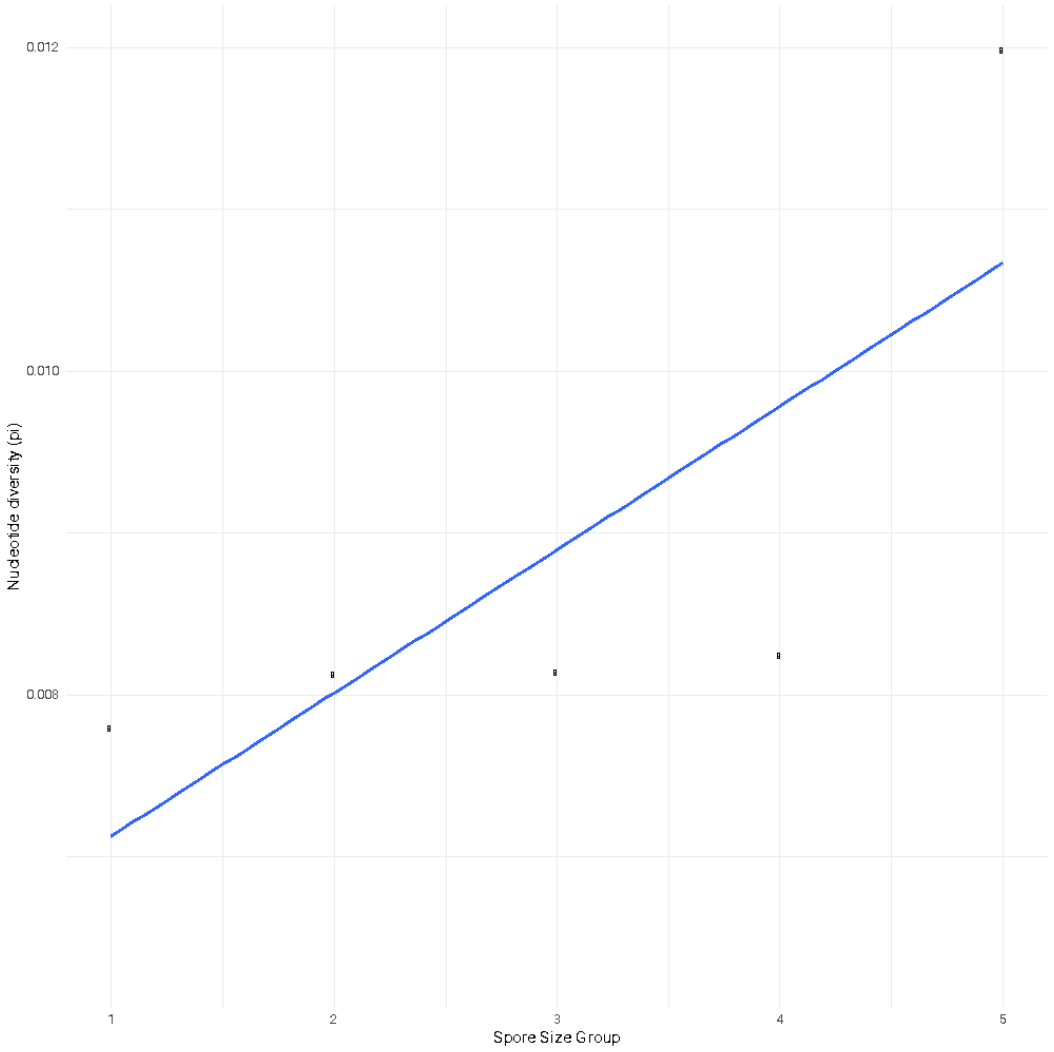


**Figure SI2.4: Linear regression for the effect of spore size and ploidy on genetic diversity (pi). Values are not reported since the relationship is driven solely by tetraploidy.**

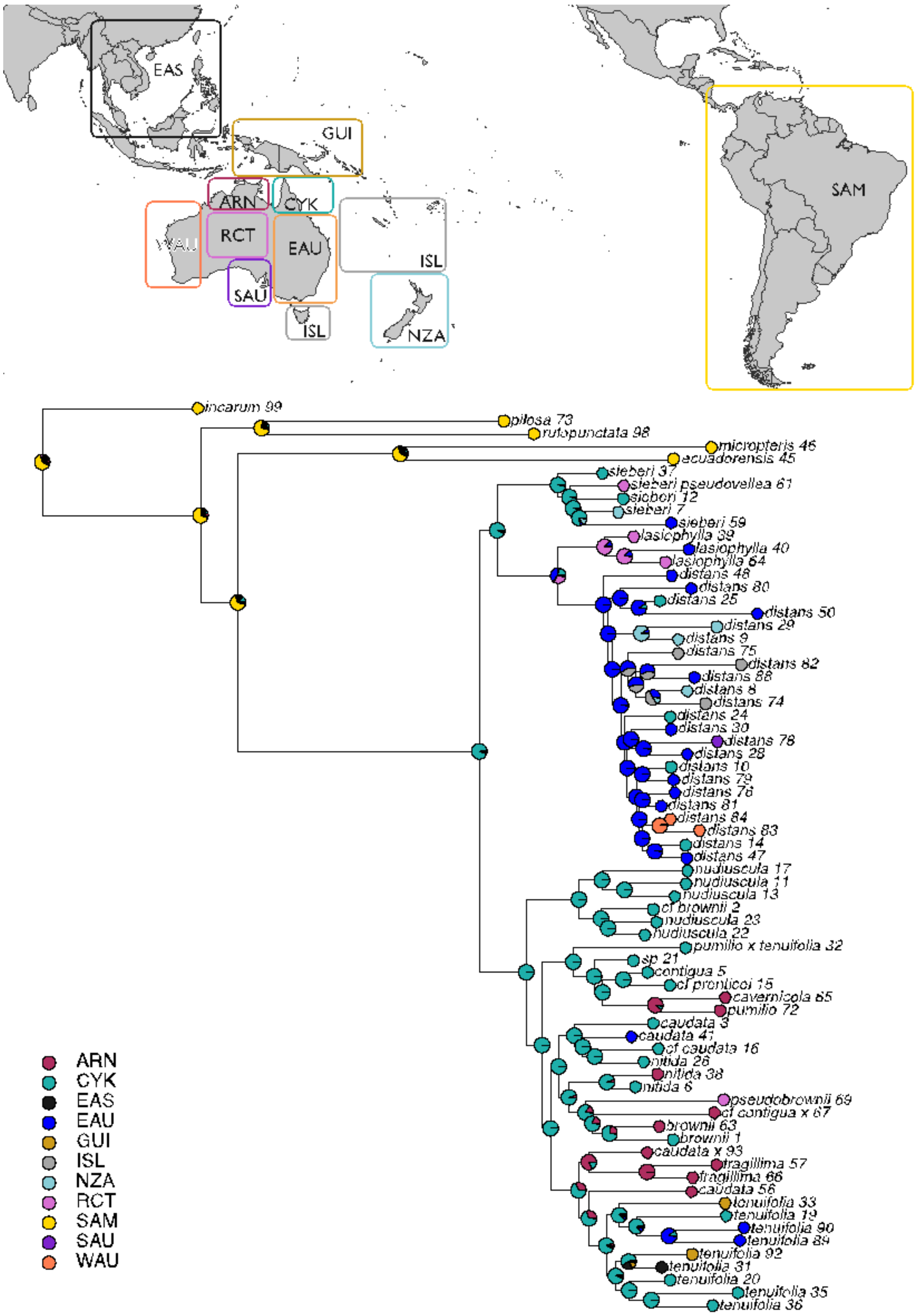


**Figure SI2.5: Ancestral state reconstructions for small geographic regions. The map above the tree shows how each of the regions is defined, as well as their colour and code.**

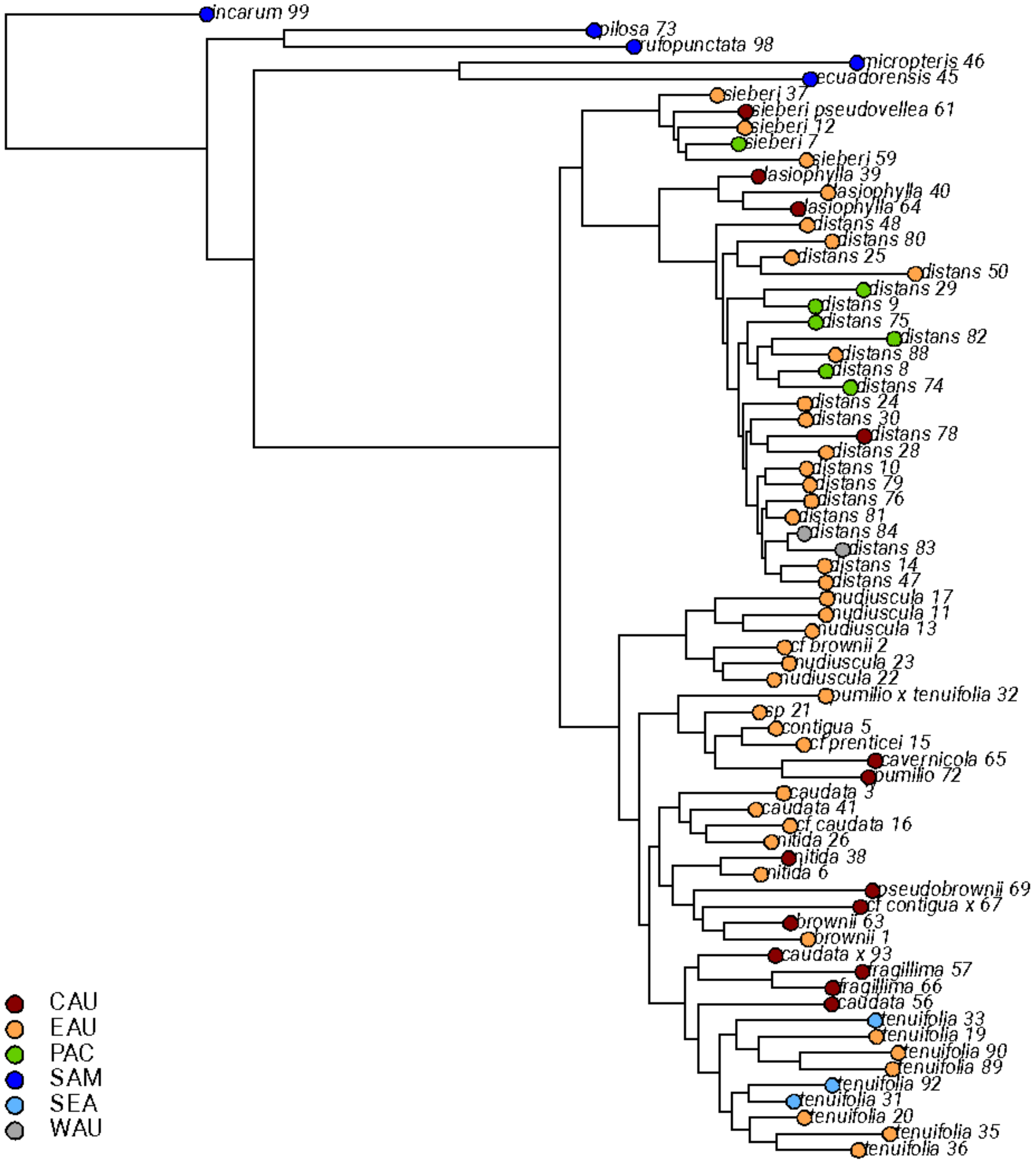


**Figure SI2.6: Summary of the stochastic character maps for shifts between large geographic regions. Each bar represents an estimated change; a total of 100 trees were sampled from the posterior and are plotted onto this phylogeny.**

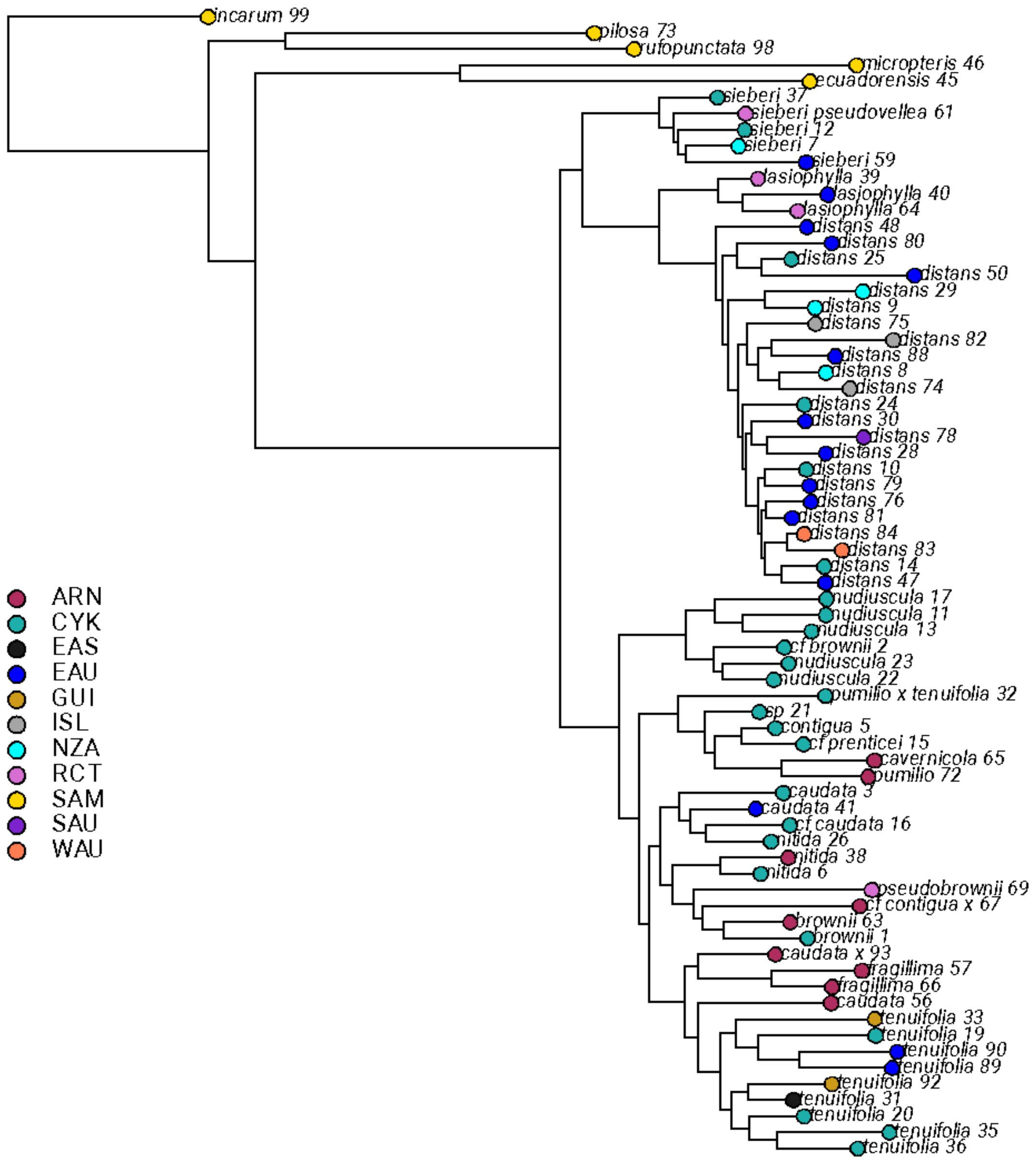


**Figure SI2.7: Summary of the stochastic character maps for shifts between small geographic regions. Each bar represents an estimated change; a total of 100 trees were sampled from the posterior and are plotted onto this phylogeny.**

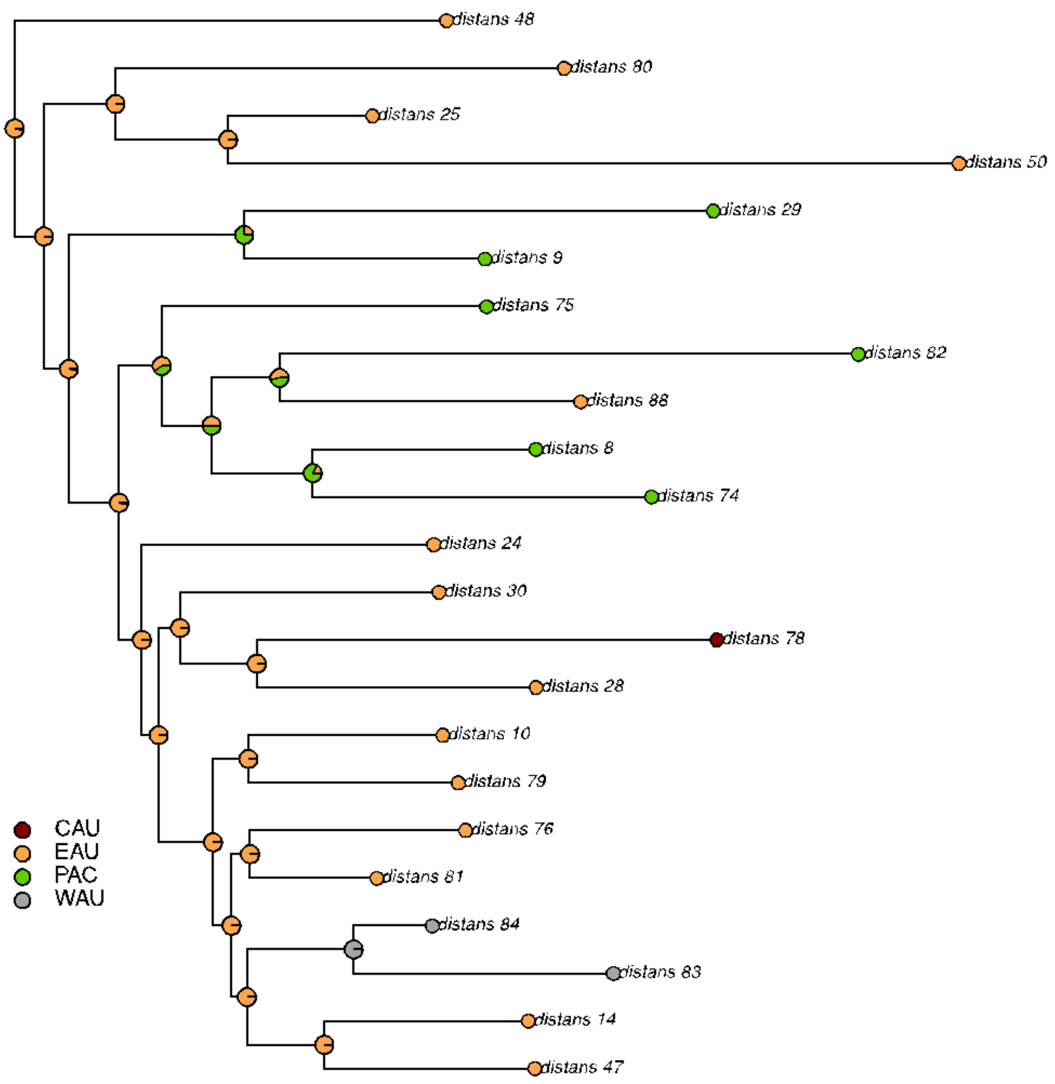


**Figure SI2.8: Ancestral state reconstructions for large geographic regions for *Cheilanthes distans* samples only.**

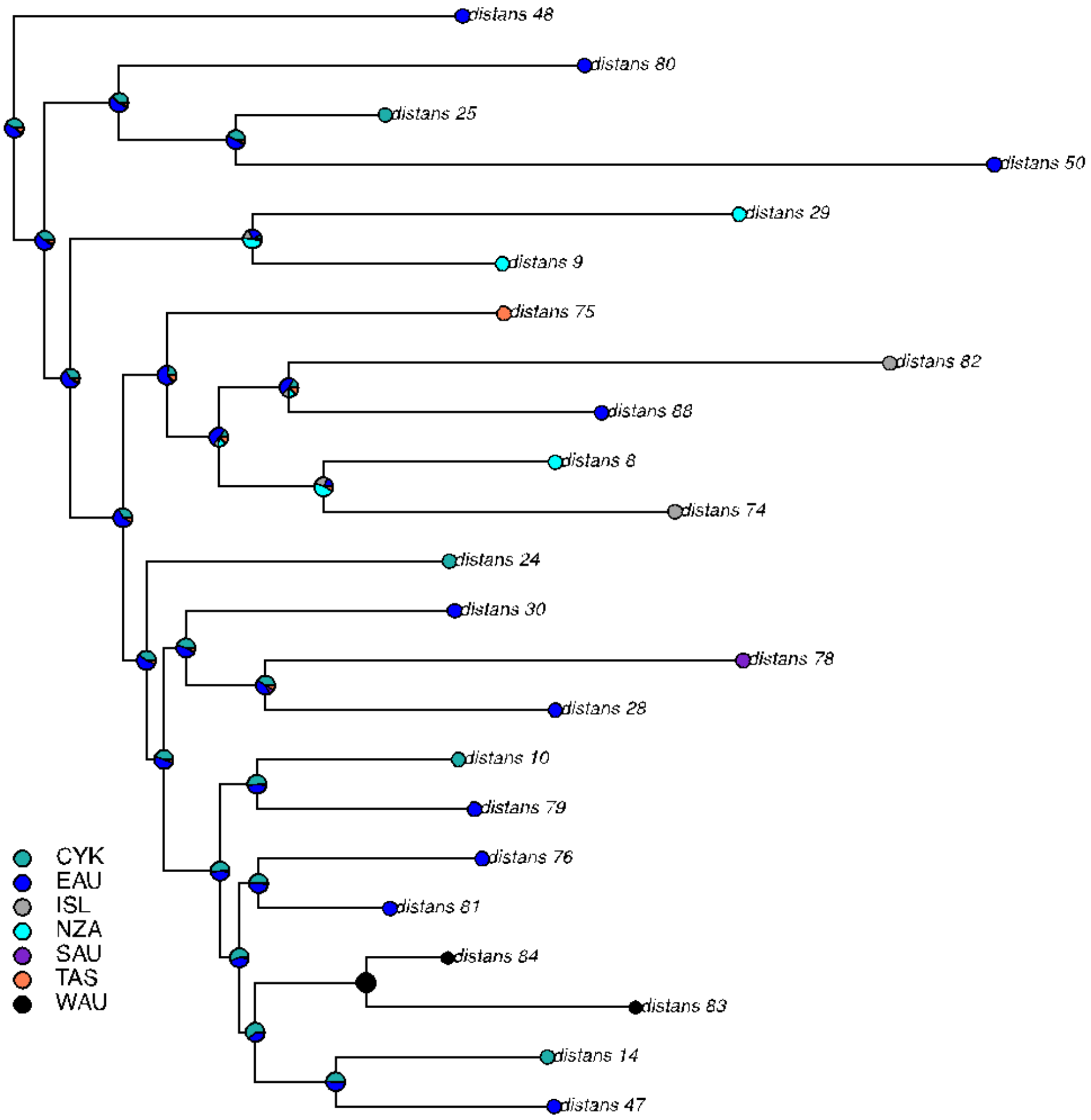


**Figure SI2.9: Ancestral state reconstructions for small geographic regions for *Cheilanthes distans* samples only.**
