## Supplemental Information 3 for "Short steps can take you far: Phylogenetic analysis of Australasian *Cheilanthes distans* reveals frequent shorter-range dispersal"

This appendix contains the phylogenetic inference results from the study. See the main text for context.

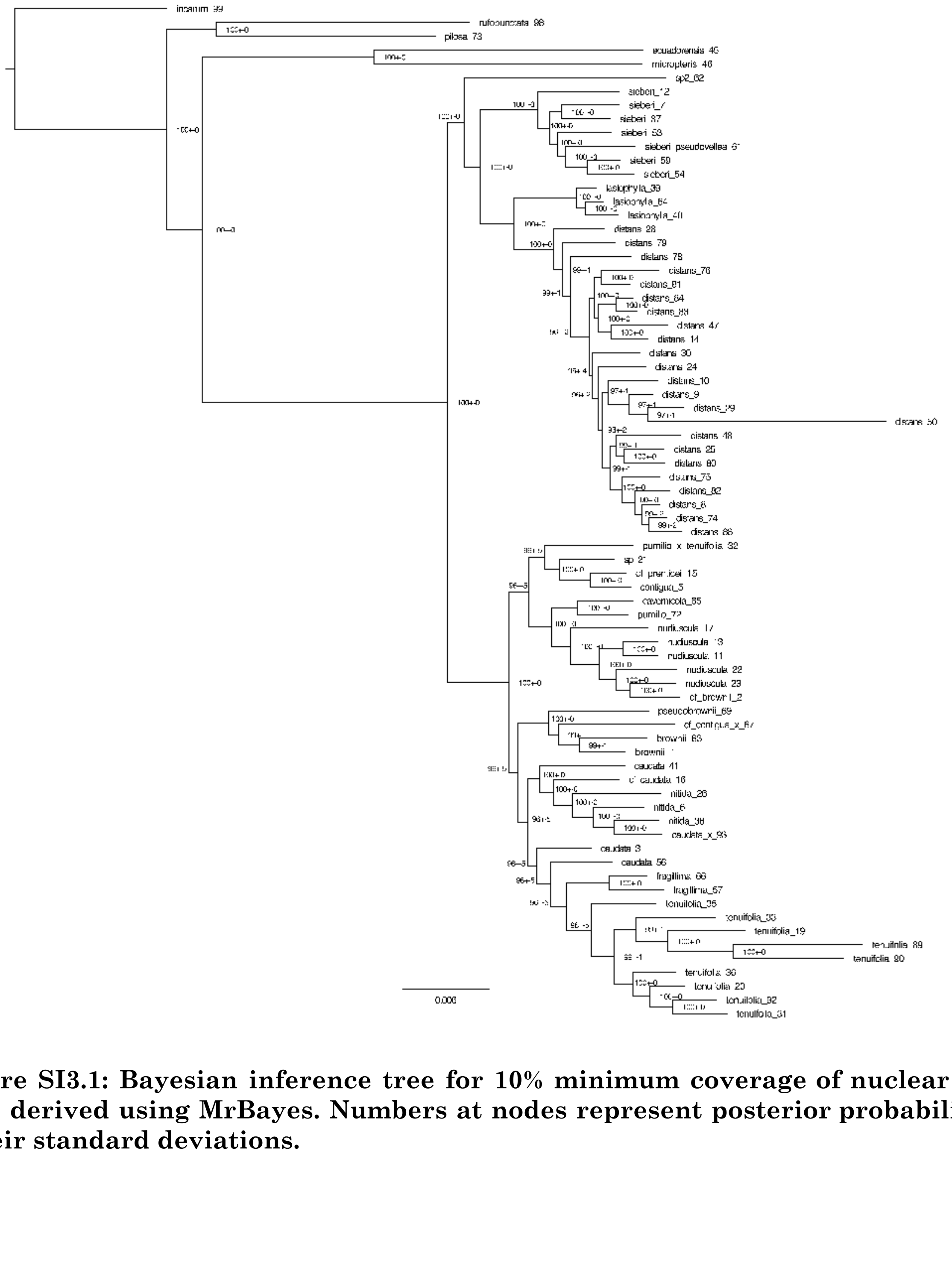


**Figure SI3.1: Bayesian inference tree for 10% minimum coverage of nuclear loci only, derived using MrBayes. Numbers at nodes represent posterior probabilities ± their standard deviations.**

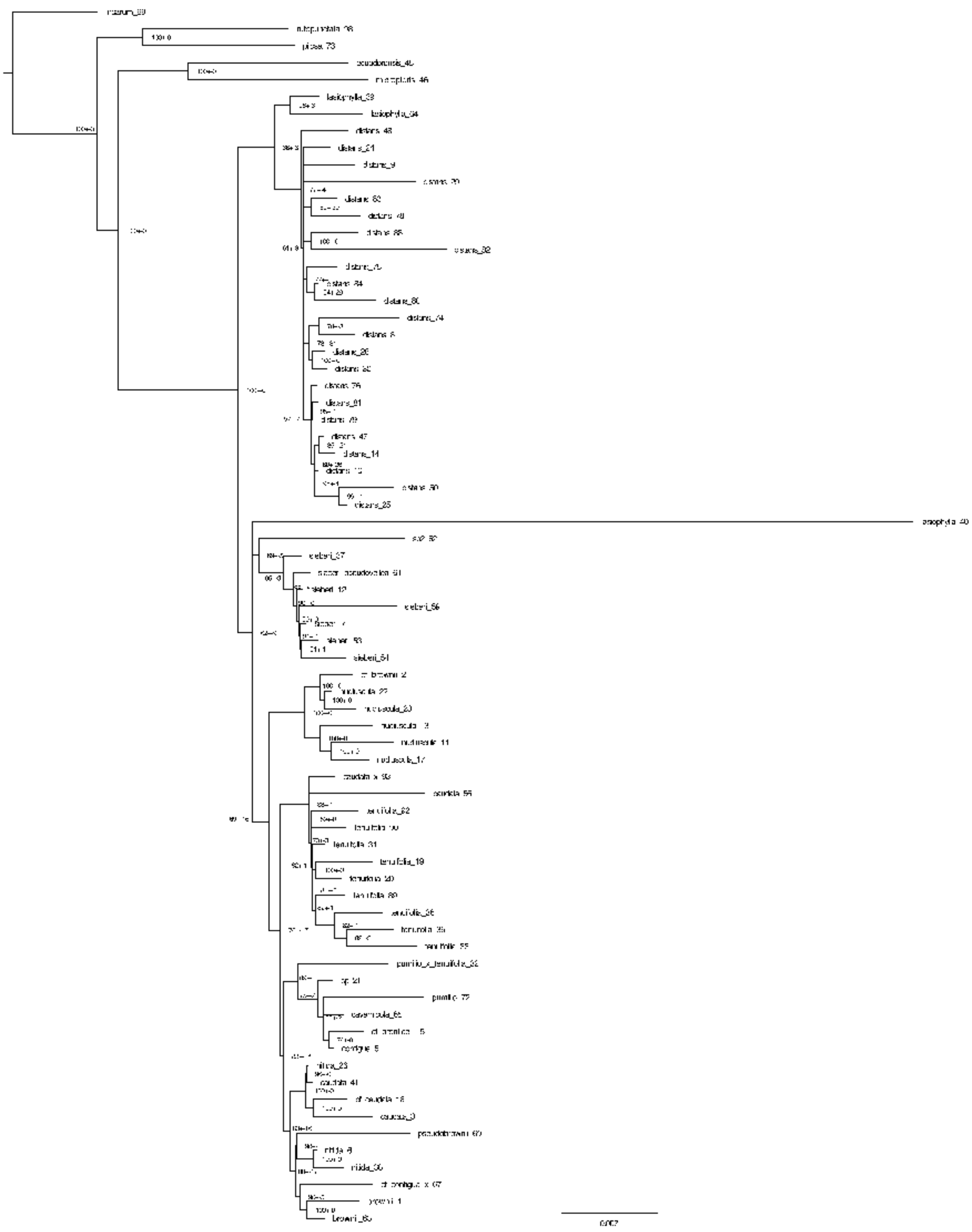

**Figure SI3.2: Bayesian inference tree for 10% minimum coverage of chloroplast loci only, derived using MrBayes. Numbers at nodes represent posterior probabilities ± their standard deviations.**

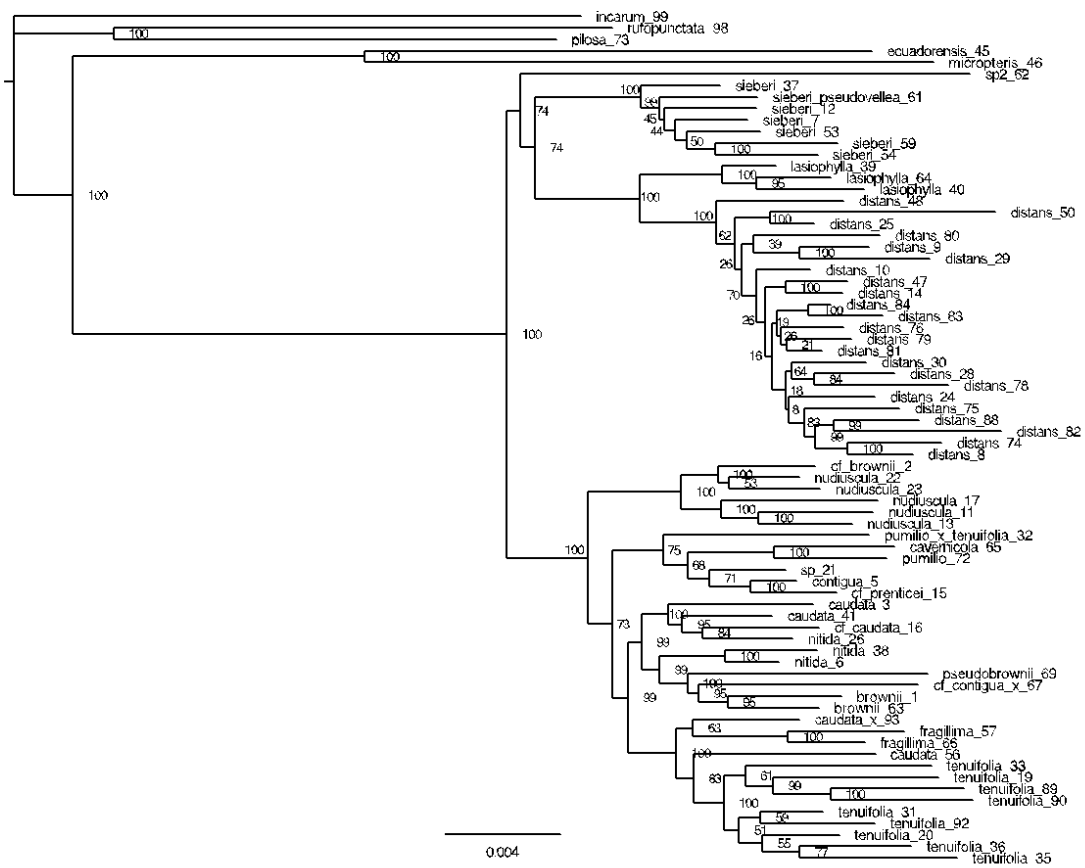

**Figure SI3.3: Maximum likelihood tree for 10% minimum coverage for combined nuclear + chloroplast datasets, estimated using RAxML. Numbers at nodes represent bootstrap values.**

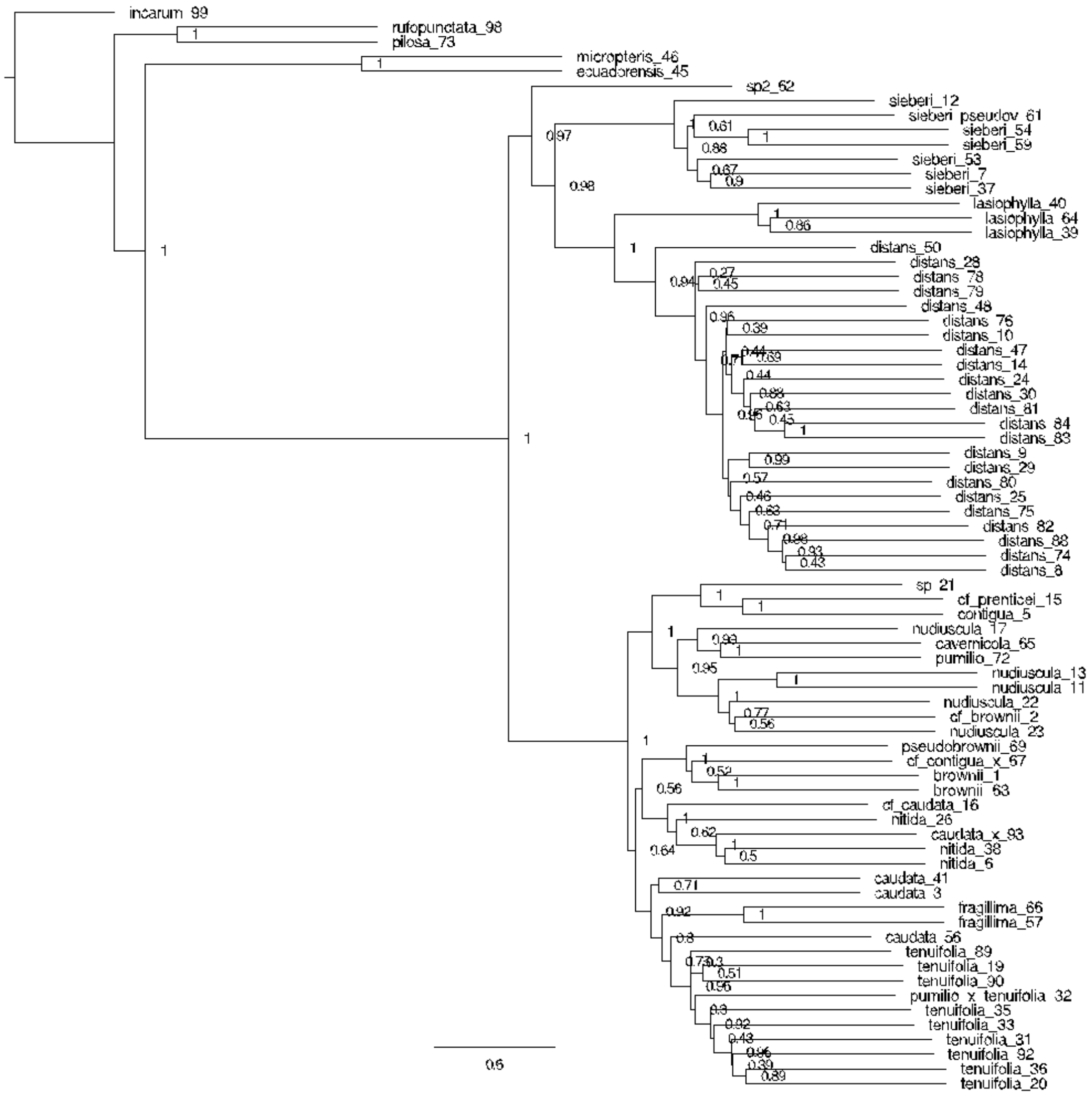


**Figure SI3.4: Coalescent tree for 10% minimum coverage for combined nuclear + chloroplast datasets, estimated using ASTRAL. Numbers at nodes represent quadripartition support values, calculated by the fraction of quartet gene trees present in the species tree.**

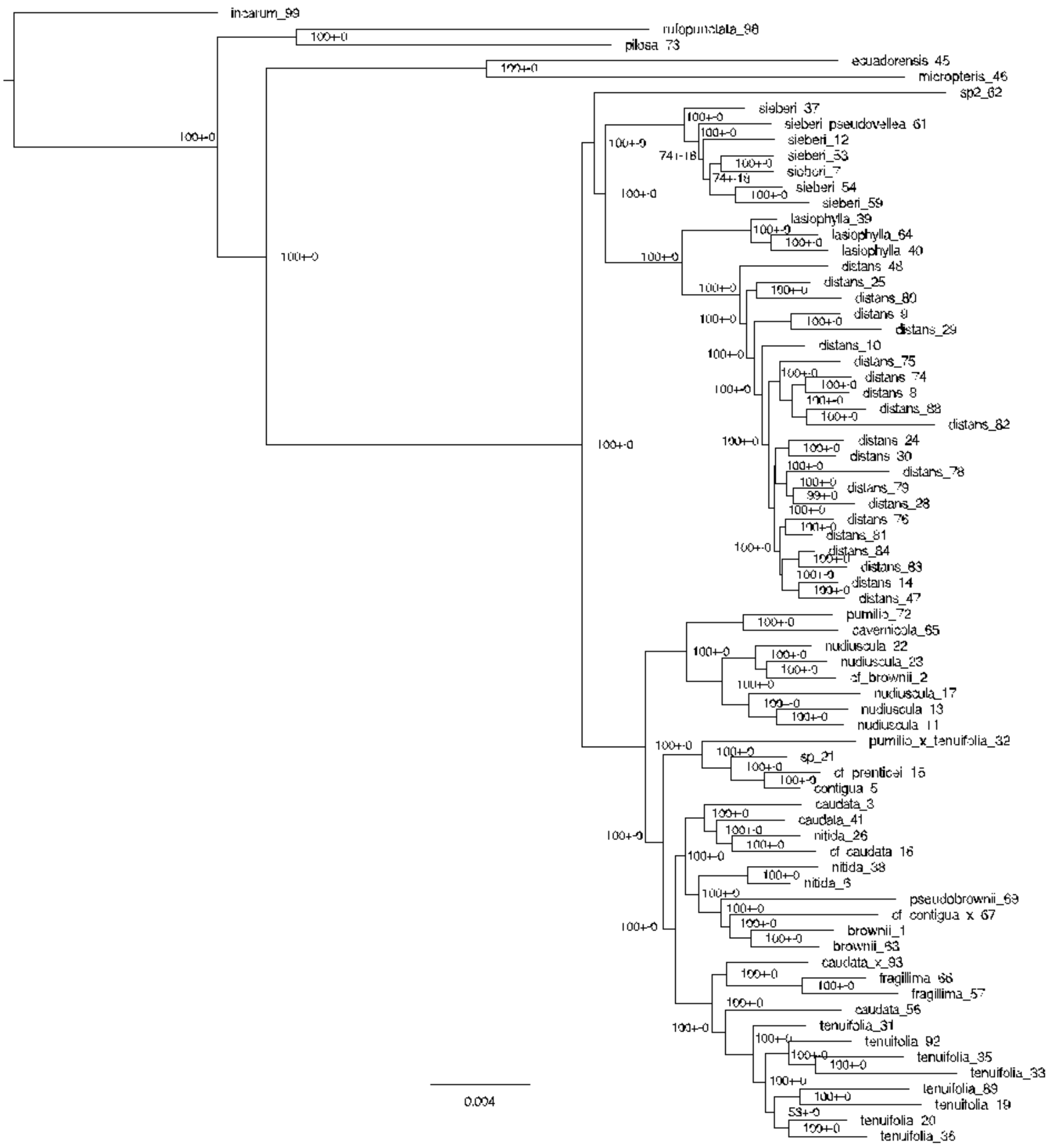


**Figure SI3.5: Bayesian inference tree for 50% minimum coverage for combined nuclear + chloroplast loci, estimated with MrBayes. Numbers at nodes represent posterior probabilities ± their standard deviations.**

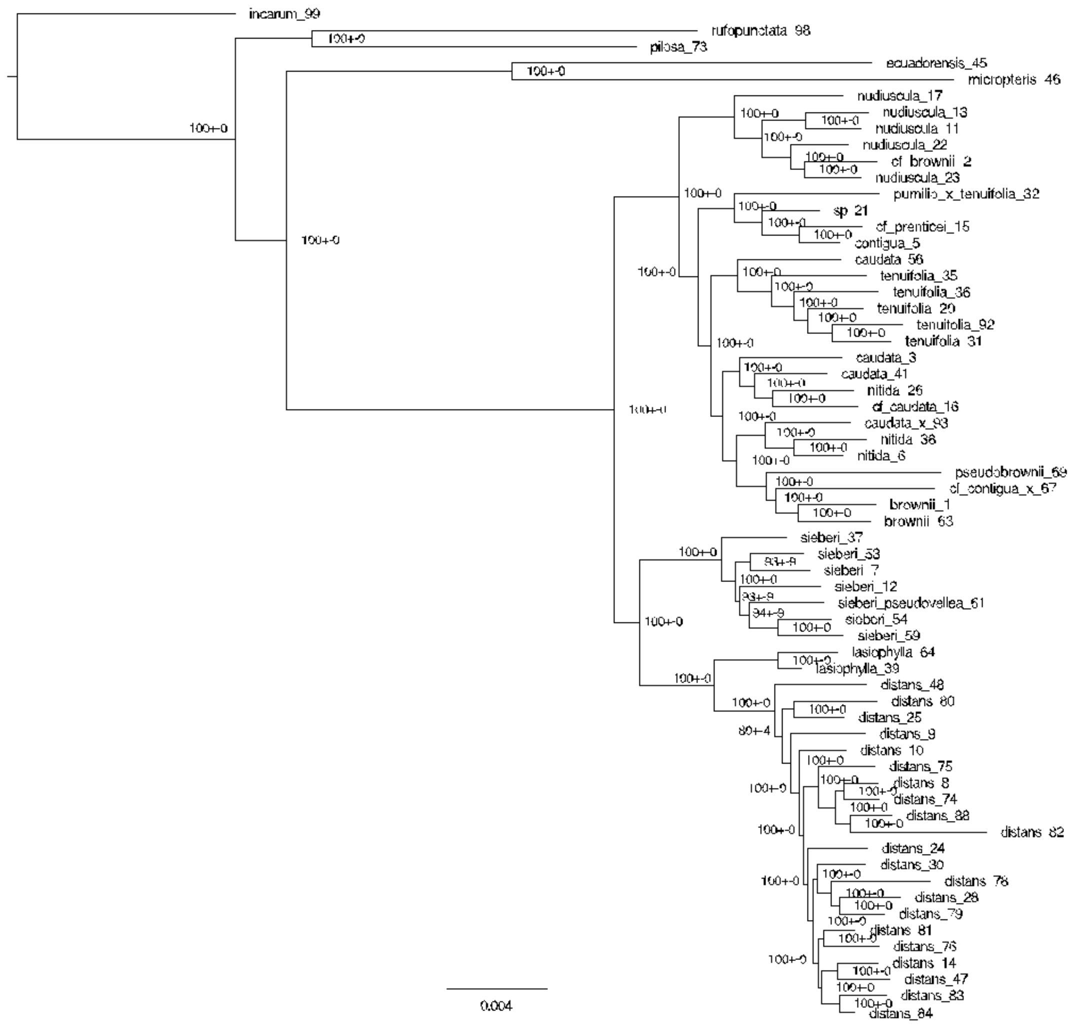


**Figure SI3.6: Bayesian inference tree for 90% minimum coverage for combined nuclear + chloroplast loci, estimated with MrBayes. Numbers at nodes represent posterior probabilities ± their standard deviations.**
